# Mycobacterial RNase E cleaves with a distinct sequence preference and controls the degradation rates of most *Mycolicibacterium smegmatis* mRNAs

**DOI:** 10.1101/2023.03.14.532454

**Authors:** Ying Zhou, Huaming Sun, Diego A. Vargas-Blanco, Maria Carla Martini, Abigail R. Rapiejko, Michael R. Chase, Samantha R. Joubran, Alexa B. Davis, Joseph P. Dainis, Jessica M. Kelly, Thomas R. Ioerger, Louis A. Roberts, Sarah M. Fortune, Scarlet S. Shell

## Abstract

The mechanisms and regulation of RNA degradation in mycobacteria have been subject to increased interest following the identification of interplay between RNA metabolism and drug resistance. Mycobacteria encode multiple ribonucleases that are predicted to participate in mRNA degradation and/or processing of stable RNAs. RNase E is an endoribonuclease hypothesized to play a major role in mRNA degradation due to its essentiality in mycobacteria and its role in mRNA degradation in gram-negative bacteria. Here, we defined the impact of RNase E on mRNA degradation rates transcriptome-wide in the non-pathogenic model *Mycolicibacterium smegmatis*. RNase E played a rate-limiting role in the degradation of at least 89% of protein-coding genes, with leadered transcripts generally being more affected by RNase E repression than leaderless transcripts. There was an apparent global slowing of transcription in response to knockdown of RNase E, suggesting that *M. smegmatis* regulates transcription in responses to changes in mRNA degradation. This compensation was incomplete, as the abundance of most transcripts increased upon RNase E knockdown. We assessed the sequence preferences for cleavage by RNase E transcriptome-wide in both *M. smegmatis* and *M. tuberculosis*, and found a consistent bias for cleavage in C-rich regions. Purified RNase E had a clear preference for cleavage immediately upstream of cytidines, distinct from the sequence preferences of RNase E in gram-negatives. We furthermore report a high-resolution map of mRNA cleavage sites in *M. tuberculosis*, which occur primarily within the RNase E-preferred sequence context, confirming RNase E as a broad contributor to *M. tuberculosis* transcriptome structure.

## INTRODUCTION

Mycobacteria are a globally important group of bacteria including the pathogen *Mycobacterium tuberculosis,* which kills over a million people each year (1), as well as numerous environmental bacteria and opportunistic pathogens. Mycobacteria are phylogenetically distant from better-studied models such as *Escherichia coli*, and consequently, numerous aspects of their fundamental biology remain poorly understood. mRNA metabolism is a critical aspect of mycobacterial biology, as regulation of gene expression facilitates adaptation to stressors both during infection and in the environment, and regulation of mRNA degradation permits energy conservation during severe stress. The roles and regulation of mycobacterial mRNA degradation enzymes remain largely undefined; however, recent reports of interplay between RNA metabolism and drug resistance have highlighted the relevance of these pathways (2–6).

The endoribonuclease RNase E is a critical component of the bulk mRNA degradation machinery in gram-negative bacteria. In *E. coli*, RNase E cleaves single-stranded mRNAs in A/U-rich regions and interacts with other RNA degradation proteins to increase the efficiency of mRNA degradation ((7–11) and reviewed in (12)). In contrast, many gram-positive bacteria such as *Bacillus subtilis* and *Staphylococcus aureus* lack RNase E completely and rely on other RNases such as RNase J and RNase Y. Mycobacteria are phylogenetically more closely related to gram-positive bacteria than gram-negatives, despite having cell envelopes that prevent gram staining. However, they encode orthologs of RNase E, and these genes are essential in both *M. tuberculosis* and the non-pathogenic model *Mycolicibacterium smegmatis* (13–15). The essentiality of RNase E suggests it may be a critical component of the bulk mRNA degradation machinery in mycobacteria. Consistent with this, mycobacterial RNase E was shown to interact with other RNases such as RNase J and PNPase (16). It was also shown to contribute to rRNA maturation (15).

We previously showed that the *M. smegmatis* transcriptome is shaped by endonucleolytic cleavage events that produce mRNA fragments with monophosphorylated 5’ ends (17). RNase E is known to produce cleavage products with monophosphorylated 5’ ends in other organisms. Taken together with the observation that the mycobacterial cleavage sites occurred preferentially in single-stranded regions, and the paucity of other candidate RNases predicted to cleave with those properties, we hypothesized that RNase E was responsible for the majority of the cleavage sites we mapped in *M. smegmatis*. However, the mycobacterial cleavage sites occurred primarily in a sequence context distinct from that reported to be cleaved by *E. coli* RNase E. Most mycobacterial mRNA cleavages occurred immediately upstream of a cytidine, with a preference for 1-2 purines immediately upstream and uridine three nt downstream of the cleavage site (RR↓**C**NU). A previous report tested the cleavage specificity of *M. tuberculosis* RNase E *in vitro*; however, the substrates used in that study did not include “RRCNU” (18).

Given the clear importance of RNase E in mycobacteria and lack of information on its role, we sought to define its function in mycobacterial mRNA metabolism. First, we used an inducible system to interrogate the effects of knockdown of *rne*, the gene encoding RNase E, in *M. smegmatis*. We found that RNase E has a rate-limiting role in degradation of most mRNAs, with a larger influence on leadered transcripts compared to leaderless transcripts. Its cleavage signature is ubiquitous across the transcriptomes of both *M. smegmatis* and *M. tuberculosis* and is distinct from that reported in gram-negative bacteria. We then used purified RNase E to confirm its cleavage specificity *in vitro*. Finally, we report a transcriptome-wide high-resolution map of major RNA cleavage sites in *M. tuberculosis,* which occur in sequence contexts corresponding to the RNase E signature. Together, our results implicate RNase E as the predominant source of 5’ monophosphorylated, cleaved mRNAs in the transcriptomes of both *M. smegmatis* and *M. tuberculosis* as well as a critical mediator of bulk mRNA degradation in these organisms.

## RESULTS

### RNase E has a global role in *M. smegmatis* mRNA degradation

Given its essentiality in mycobacteria and its broad role in mRNA degradation, we sought to determine the role of RNase E in mRNA degradation transcriptome-wide in a mycobacterial model. We therefore constructed an *M. smegmatis* strain in which we could repress transcription of *rne* (msmeg_4626), the gene encoding RNase E. Replacement of the native *rne* promoter and 5’ UTR (17) with the P766(8G) promoter and associated 5’ UTR (19) produced a strain in which anhydrotetracycline (ATc) caused a constitutively expressed reverse TetR to bind the promoter and repress *rne* transcription (Fig. 1A-B, Table 1). We hereafter refer to this as the repressible *rne* strain. Consistent with the known essentiality of *rne*, growth slowed approximately 15 hours after addition of ATc and later ceased (Fig. 1C). Construction of the repressible strain involved insertion of a hygromycin resistance gene upstream of *rne*. We therefore constructed an isogenic strain in which the hygromycin resistance gene was inserted upstream of the native copy of *rne*, hereafter referred to as the control strain (Fig. 1A).

**Figure 1.**
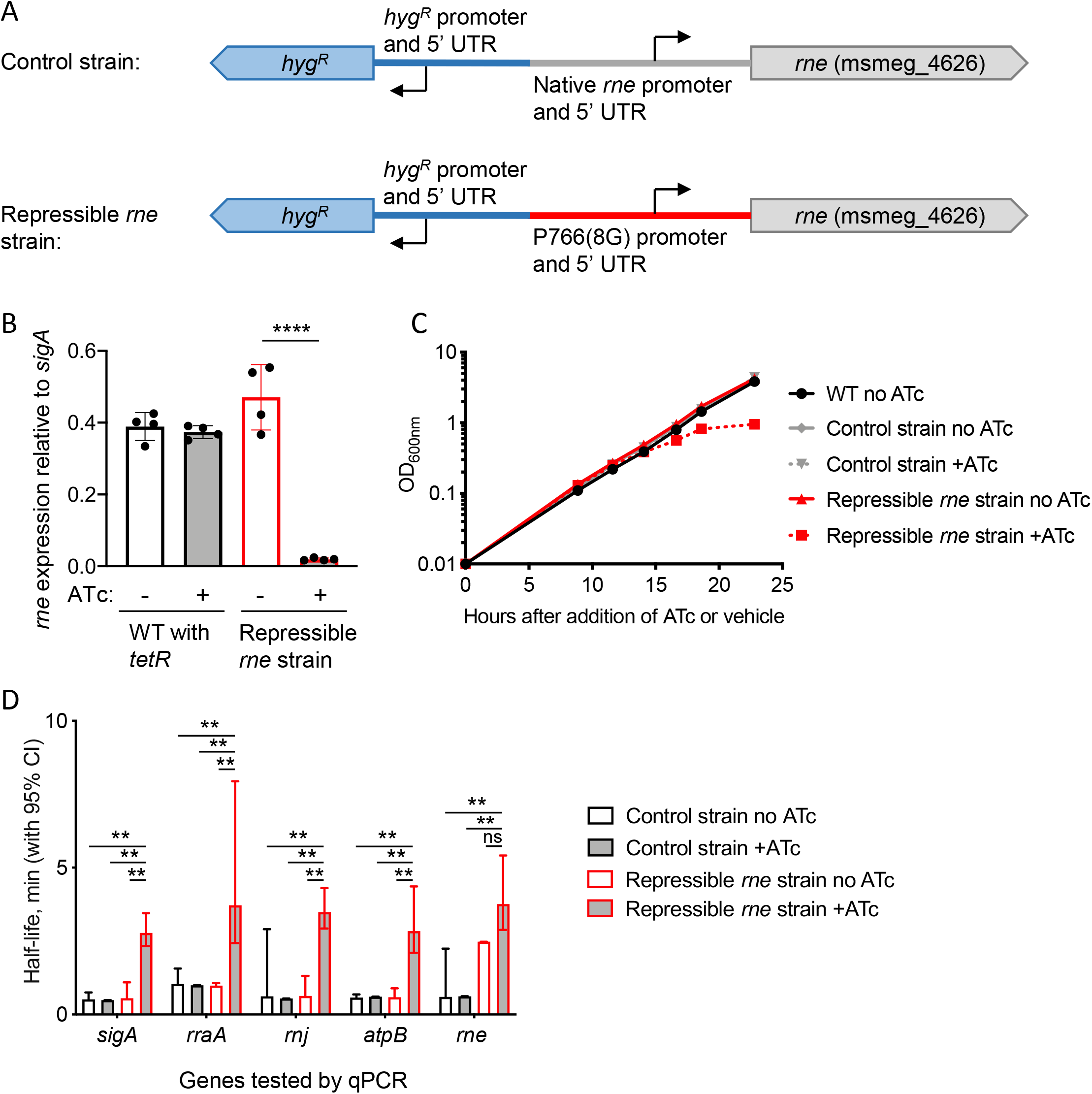
Knockdown of *rne* expression causes growth cessation and altered transcript abundance in *M. smegmatis*. **A.** Promoter replacement strategy to construct a strain in which *rne* expression is repressed by addition of ATc. **B.** *rne* transcript levels were reduced in the repressible *rne* strain following 3 hrs of exposure to ATc. **** *P* < 0.001, two-tailed t test. **C.** Growth of the repressible *rne* strain slowed approximately 15 hours after addition of ATc. **D.** Eight hours after addition of ATc or vehicle, rifampicin was added to block new transcription and mRNA levels of the indicated genes were measured at several time-points by qPCR to determine their half- lives. ** *P* < 0.01, pair-wise comparisons by linear regression.

**Table 1.** Strains and plasmids used in this study.

| Species | Strain | Plasmid | Description | Source |
| --- | --- | --- | --- | --- |
| *M. smegmatis* | mc<sup>2</sup>155 | None | Widely-used lab strain | ATCC |
| *M. smegmatis* | SS-M_0424 | pSS291: tetR38 driven by promoter ptb38, L5 integrating, kan<sup>R</sup> | mc<sup>2</sup>155 with the *hyg*<sup>R</sup> gene inserted with its own promoter 347 nt upstream of, and divergent from, the *rne* translation start site. | This study |
| *M. smegmatis* | SS-M_0418 | pSS291: tetR38 driven by promoter ptb38, L5 integrating, kan<sup>R</sup> | mc<sup>2</sup>155 in which the *rne* (msmeg_4626) promoter and UTR (nt -346 through -1 relative to the *rne* start codon) were replaced by the P766(8G) promoter and associated 5' UTR (19). Additionally, the *hyg*<sup>R</sup> gene was inserted with its own promoter upstream of, and divergent from, the P766(8G) promoter. | This study |
| *M. tuberculosis* | H37Rv | None | Widely-used lab strain | ATCC |
| *M. tuberculosis* | H37Rv Δ*rnj* | None | The *rnj* coding sequence was replaced with the *hyg*<sup>R</sup> coding sequence. | (3) |
| *E. coli* | BL21 DE3 pLysS | pSS420: pET38 expressing residues 146-824 of *M. smegmatis* RNase E with N-terminal 6XHis, 3XFLAG, TEV protease cleavage site, and 4XGly linker. |  | This work |
| *E. coli* | BL21 DE3 pLysS | pSS421: pSS420 with mutations D694R and D738R. |  | This work |

While the essentiality of *rne* could be due to its role in mRNA degradation, rRNA maturation, or both, we were specifically interested in determining the role of RNase E in mRNA metabolism. We therefore evaluated the impact of *rne* knockdown on mRNA degradation rates prior to the slowing of bacterial growth. We measured the half-lives of several mRNAs by adding rifampicin (RIF) to block transcription initiation and quantifying transcript abundance at timepoints thereafter by quantitative PCR (qPCR). The half-lives of all tested genes were lengthened upon *rne* knockdown (Fig. 1D). To determine the generalizability of this observation, we used RNAseq to measure mRNA half-lives transcriptome-wide. RNAseq libraries were constructed from RNA extracted from triplicate cultures of each strain and condition at various timepoints after the addition of RIF. qPCR was used to establish relative abundance values for a set of calibrator genes, and these were used to normalize the coverage values obtained from the RNAseq libraries as described in detail in the methods section. Libraries were made from the repressible *rne* strain following 8 hours of treatment with ATc (*rne* knockdown condition), the repressible *rne* strain in the absence of ATc, and the control strain harboring the native *rne* promoter in the presence and absence of ATc. The timepoint for analysis of the *rne* knockdown condition was carefully chosen to maximize our power to detect relevant phenotypes, but prior to the slowing of growth. We expected growth changes would themselves affect mRNA stability as has been reported by us and many others (20–27).

To identify transcripts that were direct targets of RNase E, we calculated half-lives for each gene in each condition as described in the methods section and Fig. S1-S3 (Table S1). We determined high-confidence half-lives for 1643 genes and medium-confidence half-lives for an additional 3565 genes in the *rne* knockdown condition. We were able to calculate high-confidence half-lives for 4,068 of these genes in the repressible *rne* strain in the absence of ATc as well. Half-lives were similar in comparisons between control conditions, indicating that mRNA degradation rates were not substantially affected by the presence of ATc or by replacement of the native *rne* promoter and 5’ UTR with the tet-repressible promoter (Fig. S4). In contrast, the half-lives of most genes were longer in the *rne* knockdown (Fig. 2A-B and Table S2). The half-lives of 3,622 genes increased by 2-fold or more, and an additional 78 genes had no measurable degradation in the *rne* knockdown. Together, these data are consistent with RNase E playing a rate-limiting step in the degradation of at least 89% of the transcriptome.

**Figure 2.**
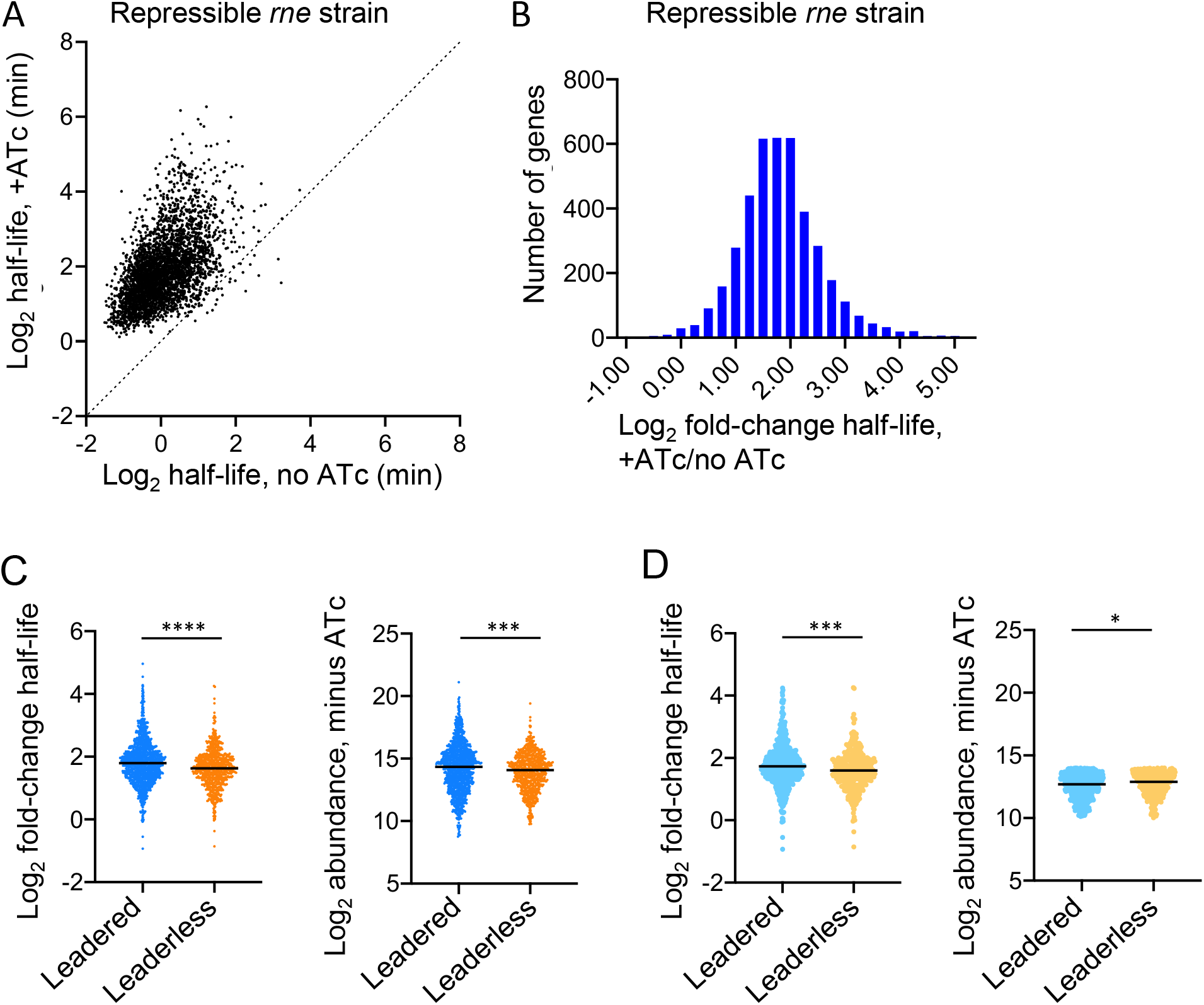
Knockdown of *rne* expression causes stabilization of most of the *M. smegmatis* transcriptome, with leadered transcripts tending to be stabilized more than leaderless transcripts. Eight hours after addition of ATc (or vehicle) to knock down (or not) *rne*, rifampicin was added to block new transcription and mRNA levels were measured transcriptome-wide at several time-points by RNAseq to determine half-lives. **A.** Dots represent genes with measurable half-lives in both conditions. **B.** The distribution of fold-change in half-life for the genes shown in A. **C.** The median fold-change in half-life upon *rne* knockdown was higher for leadered genes than for leaderless genes (left). The median abundance of leadered genes was higher prior to *rne* knockdown (right). **D.** Only genes with 10< log_2_ abundance <14 were considered, to reverse the difference in abundance trend between leadered and leaderless transcripts. The median fold-change in half-life upon *rne* knockdown was still higher for leadered genes than for leaderless genes.

While the transcripts of most genes had longer half-lives in the *rne* knockdown condition, the magnitude of the increase in half-life varied substantially among genes (Fig. 2B). To investigate the factors that influence transcript sensitivity to RNase E, we examined fold-change half-life in the *rne* knockdown as a function of other potentially relevant characteristics. There was a weak but statistically significant correlation between mRNA abundance in the control condition and fold-change in half-life upon *rne* knockdown (Fig. S5), suggesting that more abundant transcripts may be more sensitive to RNase E. Previous work has reported conflicting observations about the relationship between mRNA abundance and degradation rates in bacteria. Some studies, including one on *M. tuberculosis* and several on *E. coli*, reported inverse relationships between steady-state mRNA abundance and half-lives, such that more abundant transcripts tended to be degraded more quickly (21,24,26–30). Other studies of *E. coli* and *B. subtilis* reported that mRNA abundance and half-life were uncorrelated or weakly positively correlated (23,31,32). We found a weak but statistically significant negative correlation between mRNA abundance and half-life when *rne* was expressed at normal levels, and this correlation disappeared upon *rne* knockdown (Fig. S6).

On average, leaderless genes were less affected by *rne* knockdown than leadered genes (Fig. 2C, left). Leaderless genes also had lower median abundance than leadered genes in the control condition (Fig. 2C, right). We then considered only genes where 10<log2 abundance<14 (Fig 2D, right). Within this group, the median abundance of leaderless transcripts was slightly higher than that of leadered transcripts. Nonetheless, the leadered transcripts within this group still had a greater median increase in half-life upon *rne* knockdown than leaderless transcripts (Fig 2D, left). This suggests that the difference in response of leaderless vs leadered transcripts to *rne* knockdown cannot be explained by differences in steady-state abundance of those transcripts. Leadered transcripts may therefore be generally more sensitive to RNase E than leaderless transcripts. However, both groups included genes that were unaffected by *rne* knockdown as well as genes that were strongly affected, indicating that additional factors are likely larger drivers of RNase E sensitivity. Given that RNase E is strongly stimulated by engagement of transcript 5’ ends in *E. coli* ((33,34) and others), we considered that accessible 5’ ends might make transcripts more sensitive to RNase E. However, we did not find correlations between fold-change in half-life upon *rne* knockdown and predicted secondary structure near the 5’ ends of transcripts (Fig. S7).

### Knockdown of *rne* affects mRNA abundance through both passive and active mechanisms in *M. smegmatis*

To assess the impact of *rne* knockdown on steady-state mRNA abundance, we examined transcript abundance in the *rne* knockdown strain with and without ATc prior to transcriptional blockage with RIF. Our normalization method allowed us to measure mRNA abundance relative to total RNA abundance, in arbitrary units. As total RNA yields were similar for all strains and conditions, this roughly approximates mRNA abundance per cell, measured in arbitrary units. A large majority of genes had increased abundance upon *rne* knockdown (Table S2). We therefore could not statistically assess differential expression using a standard pipeline such as DESeq2, for which the identification of differential expressed gene relies on the assumption that mean gene expression is similar in the conditions being compared. Instead, we compared transcript abundance using Clipper, which does not rely on the specific data distributions of the two conditions (35). Of 6,922 total genes with mean read counts >0 in both conditions, 2,561 genes had increased abundance upon *rne* knockdown using cutoffs of *q* < 0.05 and fold change >= 2 (Table S3). In contrast, only 9 genes that met these criteria had decreased abundance.

There was a significant positive correlation between increase in half-life upon *rne* knockdown and increase in abundance (Spearman r = 0.3565, *p* < 0.0001; Fig. 3A). These observations are consistent with the idea that slower mRNA degradation leads to accumulation of mRNA in the cell. However, the changes in mRNA abundance were of a smaller magnitude than would be expected if transcription rates remained unchanged (compare the dashed and solid lines in Fig 3A). We therefore used the measured mRNA abundance and half-life values to calculate predicted transcription rates. A majority of genes had lower predicted transcription rates in the *rne* knockdown condition, suggesting the existence of a feedback process in which transcription is slowed to partially compensate for the longer mRNA half-lives (Fig. 3B, Table S4).

**Figure 3.**
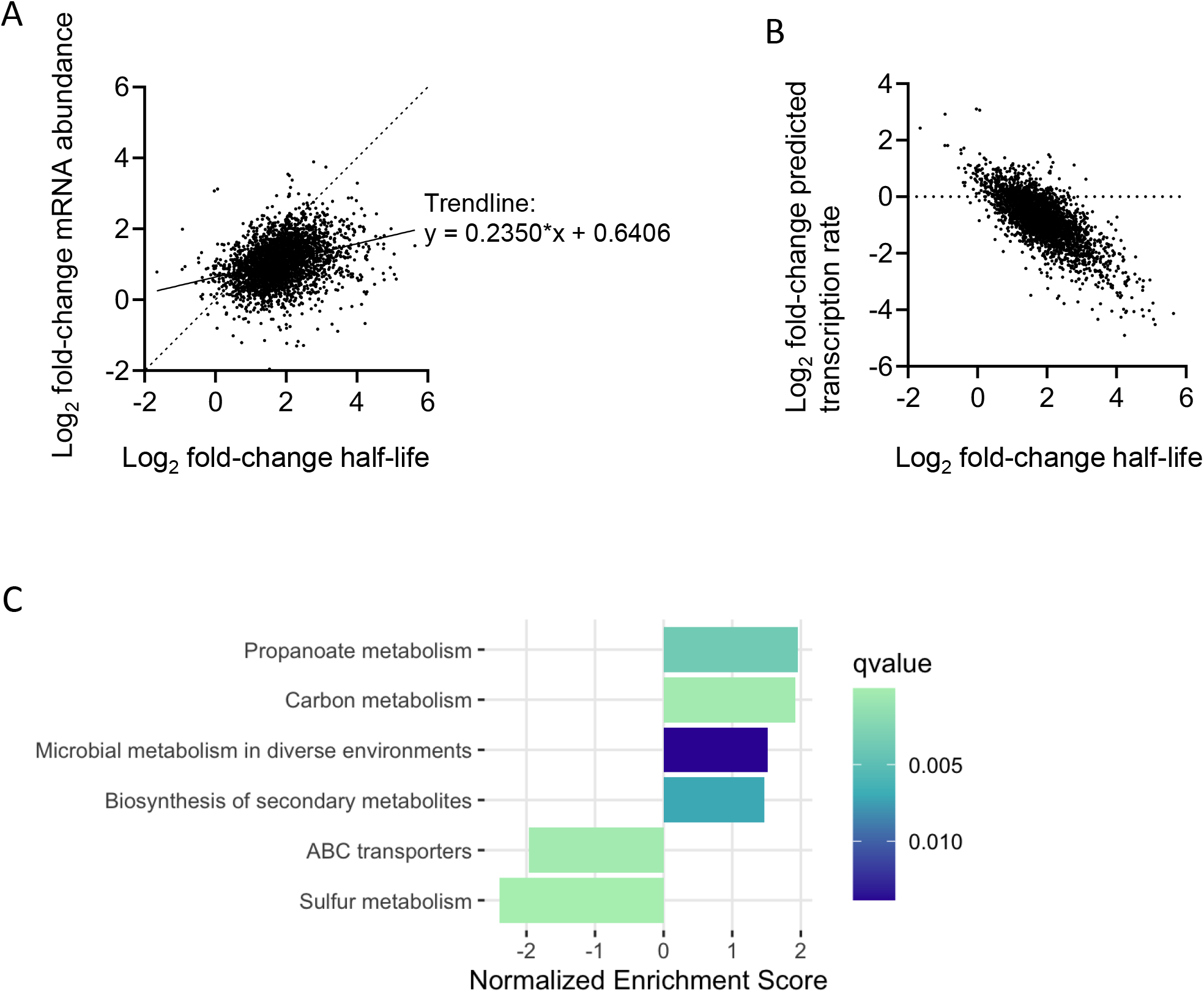
Knockdown of *rne* leads to both passive and active changes in mRNA abundance. **A.** Each dot represents a gene for which log_2_ fold change in abundance upon *rne* repression is shown as a function of log_2_ fold change in half-life. The solid line shows the linear regression fit where y = 0.2350*x + 0.6406. The dashed line shows the expected relationship between log_2_ fold change half-life and log_2_ fold change abundance if transcription rate were unchanged. **B.** Predicted transcription rates were calculated from the measured mRNA half-lives and steady-state abundance. The same genes shown in panel A are shown here. **C.** For each gene, the expected change in abundance was calculated as a function of change in half-life according to the equation in panel A. The differences between expected and observed changes in abundance were then calculated, and genes with large differences were considered more likely to be subject to active regulation. Gene Set Enrichment Analysis was performed on the observed/expected log_2_ fold change abundance, and the gene categories with statistically significant enrichment or depletion are shown. Genes in the categories with positive enrichment scores had larger than expected increases in transcript abundance, and genes in the categories with negative enrichment scores had lower that expected increases (or had decreases) in transcript abundance.

The results described above suggested that many of the transcript abundance changes caused by *rne* knockdown occurred passively due slower degradation that was only partially compensated for by globally reduced transcription. However, some genes did not follow the bulk trend. We hypothesized that the stress imposed by *rne* knockdown led to active transcriptional changes of some specific genes. To distinguish passive and active transcript abundance changes, we fit the bulk relationship between log_2_ abundance change and log_2_ half-life change by linear regression to determine predicted abundance changes as a function of change in half-life (Table S2). The difference between expected and actual abundance change reflects the extent to which a gene deviated from the bulk trend. This approach makes the assumption that most abundance changes are passive. Genes with positive differences between observed and expected abundance change had higher abundance than expected upon *rne* knockdown, while genes with negative differences had lower abundance than expected upon *rne* knockdown. To investigate the nature of the genes under active regulation, we used Gene Set Enrichment Analysis (36) to identify gene categories that were overrepresented among genes with large differences between observed and expected abundance. Genes with higher-than-expected abundance were most enriched for carbon metabolism and propanoate metabolism, while genes with lower-than-expected abundance were enriched for sulfur metabolism and ABC transporters (Fig. 3C). Transcripts for the genes encoding the RNA helicase RhlE1 (msmeg_1540) and predicted RNA binding protein KhpB (msmeg_6941) had higher-than-expected abundance, suggesting that they are transcriptionally upregulated in response to *rne* knockdown. These two proteins have reported roles as components of mycobacterial RNA degradosomes (16). It is possible that they are upregulated to partially compensate for the decrease in RNase E abundance. However, the genes encoding two other major degradosome constituents, PNPase and RNase J, did not have substantially different abundance than expected, suggesting that their abundance is not regulated in response to RNase E deficiency.

### RNase E cleavage site regions in *M. smegmatis* and *M. tuberculosis* are enriched for cytidines

Given the global role for RNase E implied by our data, we hypothesized that RNase E was the enzyme responsible for many of the mRNA cleavage events that we previously mapped (17). Those cleavage events occurred across the transcriptome at a sequence motif not previously associated with any RNase in any organism. The dominant feature of the cleavage site sequence context was a cytidine immediately downstream of the cleavage site. To assess the impact of *rne* knockdown on mRNA cleavage in *M. smegmatis*, we modified a recently published method for assessment of mRNA cleavage from standard paired-end RNAseq libraries, without construction of separate 5’-targeted libraries (37) (Fig. S8). Briefly, we calculated RNAseq coverage at each coordinate in each coding sequence, normalized to the overall abundance of the coding sequence. Coding sequences and coordinates with low coverage were filtered out. We then calculated the log_2_ ratios of coverage for each coordinate in the repressible *rne* strain in the presence vs absence of ATc, as well as for the control strain in the presence vs absence of ATc (Fig. S9 and 4A and C). If RNase E was responsible for cleavage at a particular site, we predicted that a smaller proportion of transcripts would exist in the cleaved form in the *rne* knockdown compared to the control conditions. We therefore expected coordinates that represent cleavage sites to have higher coverage in the repressible *rne* strain in the presence of ATc compared to the absence of ATc.

The distributions of log_2_ coverage ratios in the presence and absence of ATc were centered around 0 for both strains as expected (Fig. 4A and C). The distribution was broader for the *rne* knockdown strain, consistent with the expectation that RNase E levels affect the relative abundance of cleaved vs intact transcripts. RNase E both makes cleavage products and degrades cleavage products into pieces too small to be captured by our RNAseq library construction strategy; we therefore expect the effects of *rne* knockdown on the steady-state abundance of detectable cleavage products to be complex, with some cleaved RNAs decreasing in abundance and others increasing in abundance.

**Figure 4.**
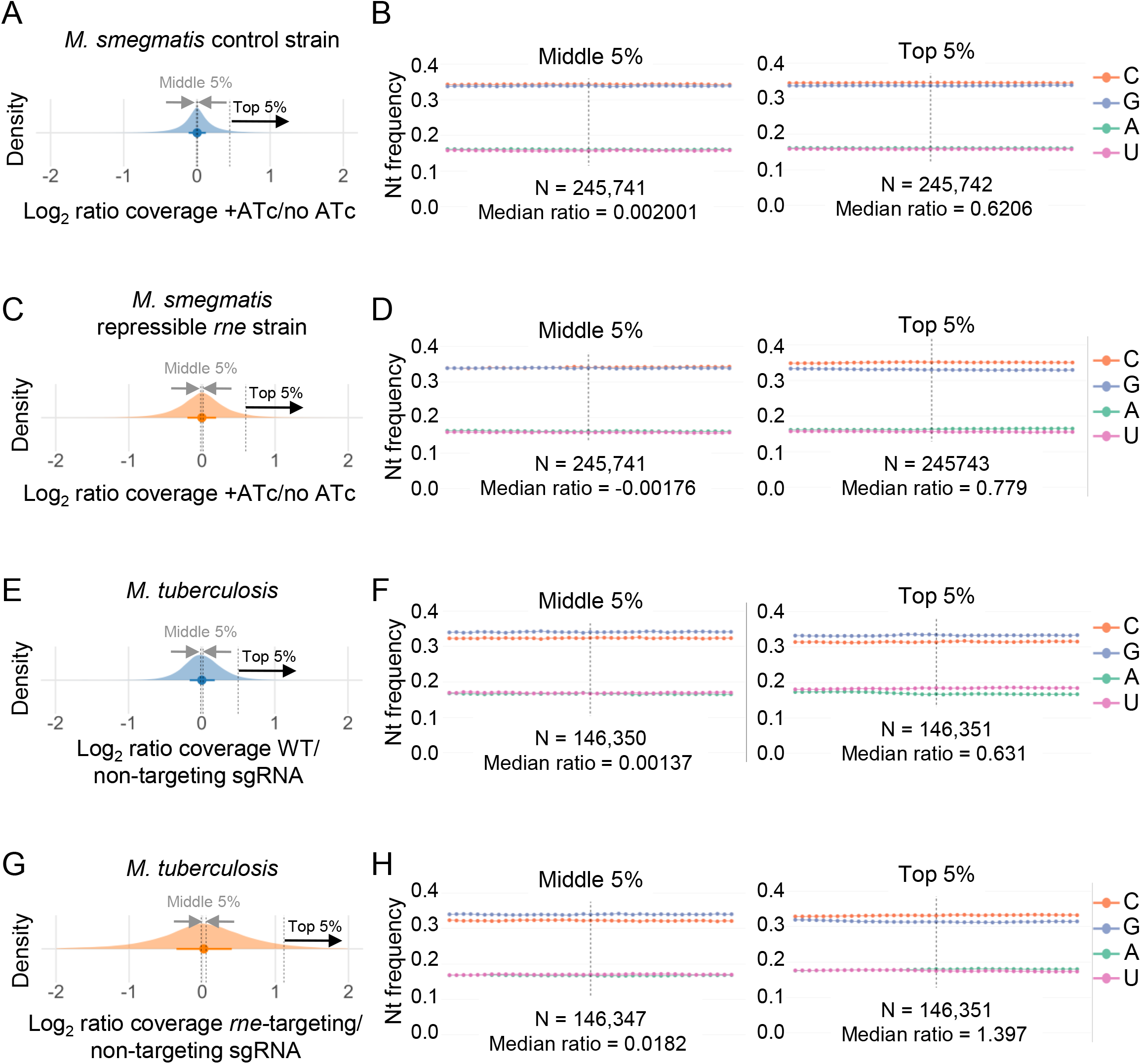
Cytidines are enriched in regions of RNase E-dependent mRNA cleavage in both *M. smegmatis* and *M. tuberculosis*. For the indicated strains and conditions, log_2_ ratios of RNAseq expression library coverage were calculated for each coordinate (distributions in panels A, C, E, G). The middle 5% and top 5% of coordinates from each distribution were selected for sequence context analysis (panels B, D, F, H). The base frequencies for the indicated coordinates and the 20 coordinates upstream and downstream are shown. **A.** Log_2_ ratios from the *M. smegmatis* control strain in the presence and absence of ATc, which is not expected to affect RNase E activity. **B.** The base frequencies corresponding to the indicated segments of panel A. **C.** Log_2_ ratios from the *M. smegmatis* repressible *rne* strain the +ATc condition (*rne* repressed) vs the no-ATc condition (*rne* expressed). Higher log_2_ ratios indicate that upon *rne* repression, there was a higher relative abundance of transcripts that were intact vs cleaved in the vicinity of the coordinate in question. **D.** The base frequencies corresponding to the indicated segments of panel B. **E.** Log_2_ ratios from two *M. tuberculosis* strains that are expected to have similar RNase E activity. **F.** The base frequencies corresponding to the indicated segments of panel E. **G.** Log_2_ ratios from an *M. tuberculosis* strain expressing an sgRNA to knock down expression of *rne* vs a strain with a non-targeting sgRNA. Higher log_2_ ratios indicate that upon *rne* repression, there was a higher relative abundance of transcripts that were intact vs cleaved in the vicinity of the coordinate in question. **H.** The base frequencies corresponding to the indicated segments of panel G.

Nonetheless, many of the coordinates at or near RNase E cleavage sites should have high log_2_ coverage ratios for the repressible *rne* strain, but this should not be true for the control strain where ATc does not affect RNase E levels. To assess the sequence context of RNase E cleavage sites, we therefore compared the sequence contexts of coordinates with the highest 5% of log_2_ ratios and the middle 5% of log_2_ ratios for both strains. The relative frequencies of each base were equivalent for the middle 5% and highest 5% of coordinates for the control strain (Fig. 4B), with G and C having similar frequencies that were much higher than A and U, as expected for an organism with a genomic GC content of ∼65%. The same was true for the middle 5% of coordinates for the *rne* knockdown strain (Fig. 4D). In contrast, the highest 5% of coordinates in the repressible *rne* strain showed a clear enrichment for cytidines (Fig. 4D). This is consistent with the hypothesis suggested by our previous work that RNase E has a preference for cleaving near cytidines (17). The enrichment for cytidines appears modest compared to our previous finding that >90% of mapped cleavage events were immediately upstream of cytidines (cytidine at the +1 position). However, this modest enrichment is consistent with the nature of the method. At any given endonucleolytic cleavage site, coordinates at the -1 position are expected to have equally high log_2_ ratios as coordinates at the +1 position, but only the +1 position shows a preference for cytidines. Furthermore, nearby coordinates (eg, -2, -3, -4, +2, +3, +4) are also likely to have relatively high log_2_ ratios. Cytidines were not enriched at any position besides +1 in our previously mapped cleavage sites, and in several of those positions there was reduced presence of cytidines (17). Adding to the complexity of the interpretation of these data, RNAseq expression library coverage is typically bumpy, with stochastic factors leading to variability in coverage among adjacent nt. The coordinates with the highest 5% of log_2_ ratios are therefore likely to include many -1 and +1 positions of cleavage sites, but also many other coordinates that are in the general vicinities of cleavage sites. The observed modest enrichment of cytidines in Fig. 4D is broadly consistent with the averaging of the previously observed sequence preferences in the vicinity of cleavage sites.

RNAseq data have been previously published for *M. tuberculosis* with *rne* knockdown (16). We therefore applied the method described above to investigate the extent to which *M. tuberculosis* RNase E preferentially cleaves cytidine-rich regions. As a control, we compared normalized RNAseq coverage from WT H37Rv to a strain expressing a non-targeting CRISPRi sgRNA (Fig. 4E). The coordinates with the middle 5% and top 5% log_2_ ratios had similar sequence contexts, which differed from the *M. smegmatis* data in having a greater proportion of guanosines than cytosines (Fig. 4F). This is consistent with differences in the overall nucleotide usage in the two organisms; *M. tuberculosis* coding sequences contain more guanosines than cytidines, while *M. smegmatis* coding sequences have roughly equal usage of guanosines and cytidines (Fig. S10). To assess the impact of RNase E, we compared a strain containing an *rne*-targeting CRISPRi sgRNA to the non-targeting sgRNA strain (Fig. 4G). The coordinates with the middle 5% of log_2_ ratios had base frequencies similar to the control comparison, but the coordinates with the highest 5% of log_2_ ratios showed a higher frequency of cytidines compared to guanosines (Fig. 4H). The preference for RNase E to cleave cytidine-rich regions is therefore conserved in M*. tuberculosis*.

### *M. smegmatis* RNase E cleaves immediately 5’ of cytidines

To assess RNase E’s cleavage site sequence preference with higher resolution, we performed two additional analyses. First, we used 5’ RACE to qualitatively compare the abundance of 5’ ends arising from a putative RNase E cleavage event in the rRNA precursor (Fig. S11A). We mapped a 5’ end in the spacer region between the 16S and 23S rRNAs resulting from cleavage at the sequence UG↓CU (Fig. S11A). Consistent with the idea that RNase E is responsible for cleaving this site, the band corresponding to the 5’ end produced by the cleavage event was fainter in the *rne* knockdown (Fig. S11B). This is consistent with a previously reported role for RNase E in cleaving near this location (15), although the method used in that report did not permit precise identification of the 5’ end as we did here.

Next, we overexpressed and purified *M. smegmatis* RNase E in *E. coli* to test its cleavage specificity in vitro. This recombinant RNase E lacked part of the predicted N-terminal scaffold domain (deletion of residues 2-145) and most of the predicted C-terminal scaffold domain (deletion of residues 825-1037), similar to RNase E variants used for in vitro work in many reports (including (18,34,38)). Our RNase E also had N-terminal 6x-His and FLAG epitope tags to facilitate purification. A variant containing the predicted catalytic site mutations D694R and D738R was purified to use as a catalytically dead control (34). The purified proteins were incubated with an in vitro-transcribed substrate that contained a duplex region and a single-stranded region (Fig. 5B and S12). Bands that appeared only in the digest with the catalytically active enzyme were subject to 5’ and 3’ RACE to map the cleavage site locations. We mapped four distinct cleavage sites, all in the single-stranded portion of the substrate (Fig. 5A-B). Two were at positions where we previously mapped cleavage sites in vivo (17), and all four occurred at the sequence motif RN↓CNU. These data confirm the propensity of RNase E to cleave single-stranded RNAs at phosphodiester bonds 5’ of cytidines.

**Figure 5.**
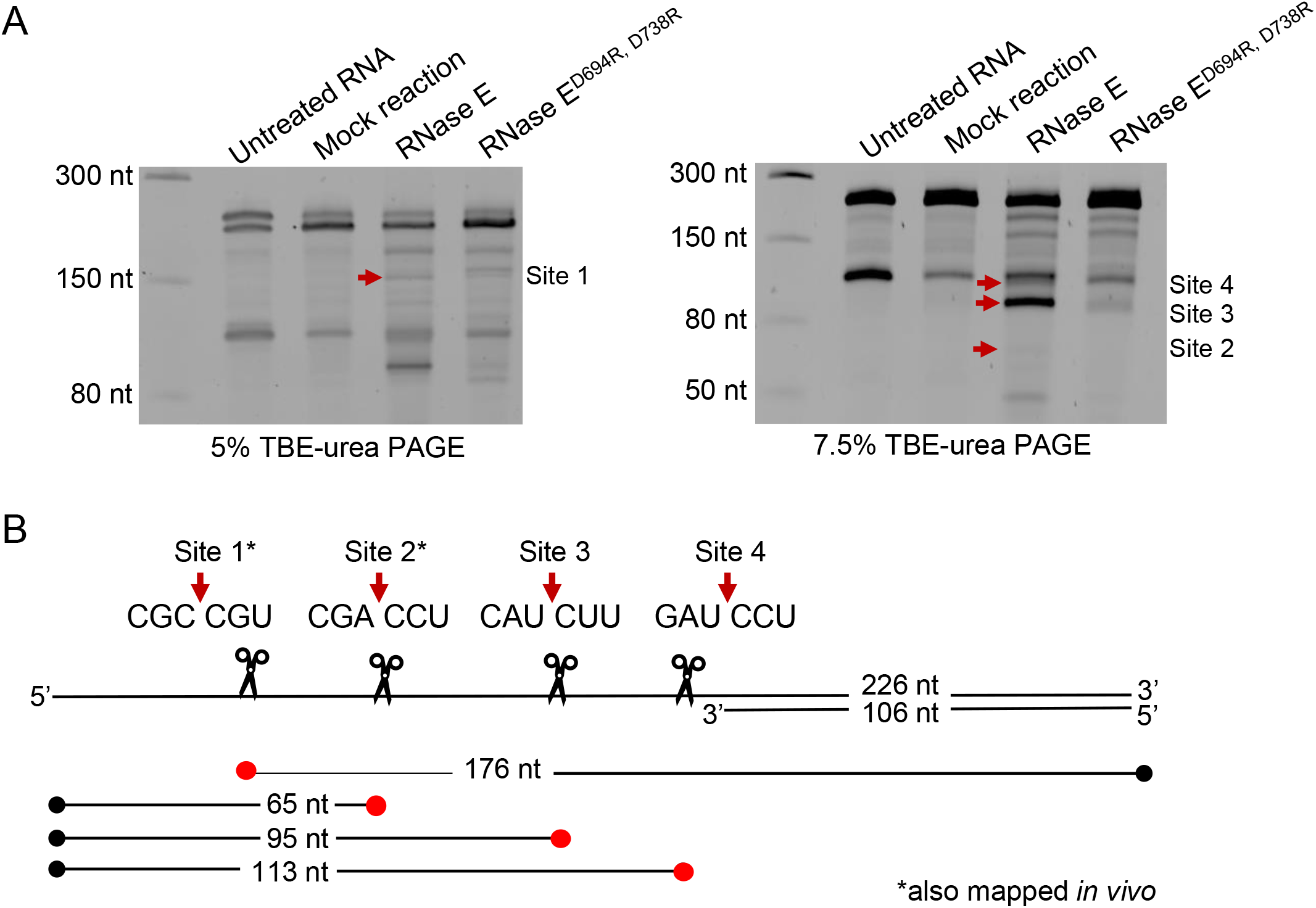
RNase E cleaves 5’ of cytidines in vitro. **A.** SYBR-gold-stained TBE-UREA gels revealing cleavage of the RNA substrate shown in B upon incubation for 1 hr with purified, recombinant *M. smegmatis* RNase E catalytic domain (residues 146-824, with an N-terminal FLAG-his tag). The D694R, D738R mutant is predicted to be catalytically dead. **B.** Schematic (not to scale) of the partial duplex substrate used in the panel A cleavage reactions. Red arrows indicate RNase E cleavage sites mapped by 5’ or 3’ RACE on cleavage products extracted from the gels shown in panel A. The thin lines below indicate the sizes of the extracted cleavage products, with red dots indicating the ends that were mapped by RACE.

### The *M. tuberculosis* transcriptome is shaped by mRNA cleavage immediately upstream of cytidines

We previously mapped *M. smegmatis* RNA cleavage sites in vivo by differential ligation (17). These data are complementary to the cleavage site analysis described above; they do not give information on the RNase responsible, but they give single-nt resolution. To determine the extent to which cleavage patterns were similar in the pathogen *M. tuberculosis*, we applied the differential ligation approach. This well-validated method distinguishes between mRNA cleavage sites and primary 5’ ends produced from transcription initiation (TSSs) based on their different chemical properties (39,40). We identified 2,983 cleavage sites with high confidence (Table S5A), using a filter that required the cleaved 5’ ends to pass an abundance threshold relative to nearby expression library coverage as we did previously for *M. smegmatis*. The TSSs mapped with this approach have been reported elsewhere (40). However, the relationships between the TSSs and genes, as well as operon predictions based on TSS locations, were not previously published and are therefore reported here in Table S5B-H.

RNA cleavage in *M. tuberculosis* occurred at a sequence motif very similar to that observed in *M. smegmatis*, with a strong bias for cleavage 5’ of cytidines (88% of high-confidence cleavage sites) and a weak bias for cleavage 3’ of purines (Fig. 6A and (17)). Given the multiple lines of evidence shown above indicating that RNase E cleaves in this sequence context, we hypothesize that RNase E was responsible for most of the mapped *M. tuberculosis* cleavage sites. Analysis of the predicted secondary structure in the vicinity of cleavage sites revealed that cleavage occurred in regions more likely to be single-stranded (Fig. 6B), consistent with expectations for RNase E (reviewed in (12)). We then removed one of the abundance filters used in the 5’ end data analysis pipeline to capture a greater number of putative cleavage sites (Table S5I). Analysis of the sequence context of this expanded cleavage site list revealed a similar preference for cleavage immediately upstream of cytidines (85% of the 5’ ends in the dataset), with a similar but weaker signal for sequence preferences at other positions surrounding the cleavage site (Fig. S13).

**Figure 6.**
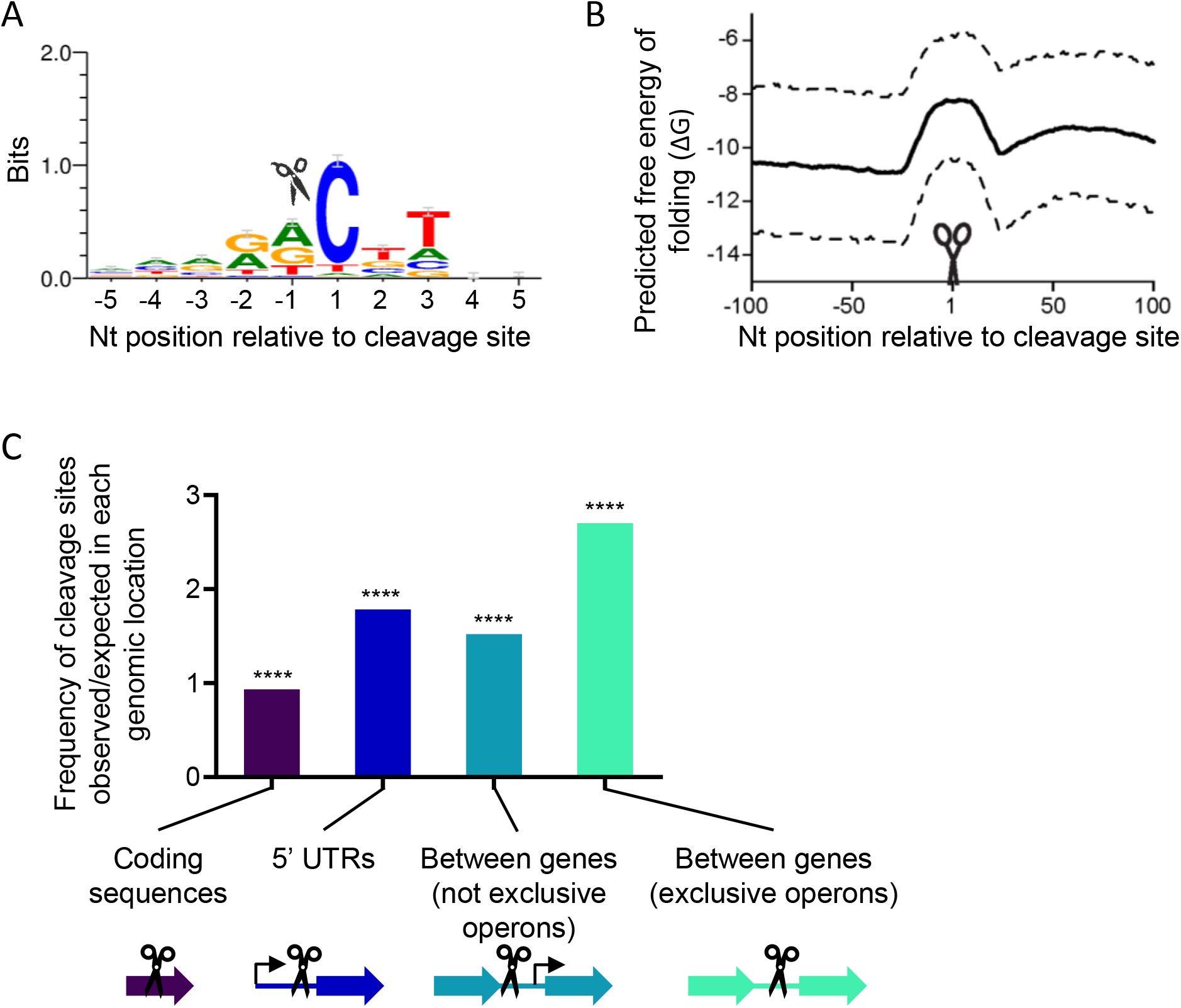
A transcriptome-wide mRNA cleavage site map in *M. tuberculosis* reveals sequence and secondary structure preferences consistent with RNase E, and greater cleavage site frequency in 5’ UTRs and intergenic regions. **A.** Weblogo (3.7.4) generated from the complete set of mapped *M. tuberculosis* cleavage sites aligned by cleavage site position. Cleavage occurs between positions -1 and 1 as indicated by the scissor icon. **B.** RNA cleavage typically occurs within regions of lower secondary structure. The minimum free energy secondary structure was predicted for sliding 39 nt windows across 200 nt of sequence spanning each RNA cleavage site. For each coordinate, the mean (solid line; interquartile range, dashed lines) predicted free energy (ΔG) of secondary structure formation of all 2,983 cleaved RNAs was determined. **C.** The frequencies of RNA cleavage sites in coding sequences, 5’ UTRs, and between adjacent genes were compared to the expected frequencies based on how much of the expressed genome comprises each of these features. 5’ UTRs were included only if the next upstream gene was on the opposite strand. Regions between genes were separated according to whether or not the gene pair was predicted to be transcribed in an exclusively operonic (polycistronic) fashion. Only regions between genes on the same strand were considered. ****, *p* < 0.0001 by binomial test comparing the observed vs expected frequencies.

MazF was reported to also cleave near cytidines (41), but it produces 5’ hydroxyls rather than 5’ monophosphates and its cleavage products are therefore not captured by our methodology. However, RNase J is predicted to cleave single-stranded RNAs and produce 5’ monophosphates. To determine if RNase J contributed to the mapped cleavage sites in *M. tuberculosis*, we compared the abundance of cleavage-site-derived 5’ ends in a WT strain and an RNase J deletion strain (Fig. S14) (3). Most of the cleaved 5’ ends had similar abundance in the two strains, consistent with the hypothesis that most of them are not produced by RNase J.

### mRNA cleavage sites are disproportionately located in 5’ UTRs and intergenic regions in *M. tuberculosis*

To further investigate the contributors to RNA cleavage site selection in *M. tuberculosis*, we examined the frequencies of cleavage sites in coding sequences, 5’ UTRs, and between adjacent genes encoded on the same strand. In each case we assessed enrichment or depletion by comparing the observed number of cleavage sites to the number expected if cleavage was equally likely to occur in those various locations. Cleavage sites were present at less than the expected frequency in coding sequences, and at greater than the expected frequency in 5’ UTRs and intergenic regions (Fig. 6C). This pattern is similar to what we previously observed in *M. smegmatis* (17). It could be the result of differential occurrence of cleavage in these locations, or could be the result of cleavage in non-coding sequences being more likely to result in products stable enough to be detected. Cleavage events that trigger very rapid degradation would be unlikely to be detected by our methodology. Interestingly, the greatest enrichment for mapped cleavage sites occurred between genes that were transcribed exclusively as polycistrons (Fig. 6C). In some cases there was differential abundance of the transcripts corresponding to genes upstream and downstream of the cleavage site (Table S6), suggesting that cleavage could lead to differential stability of segments of transcripts as has been reported in some other bacterial operons (42–48). Consistent with this idea, we found that two pairs of polycistronic *M. smegmatis* genes with intervening cleavage sites had differential stabilities upstream and downstream of the cleavage sites (Fig. S15).

## DISCUSSION

Here we used a combination of approaches to define the role of RNase E in mycobacterial mRNA degradation and identify its targets. The dramatic effect of *rne* knockdown on mRNA degradation rates in *M. smegmatis* is consistent with the essentiality of this enzyme in mycobacteria; it appears to play a rate-limiting step in degradation of the transcripts of almost 90% of genes. There was variability in the extent to which transcripts were stabilized upon *rne* knockdown, suggesting that while RNase E likely contributes to degradation of most mRNAs, other RNases may contribute differentially across the transcriptome. For example, the essential exoribonuclease PNPase could conceivably be the major degradation factor for those genes that were minimally affected by *rne* knockdown. An alternative explanation is that some mRNAs may be exquisitely susceptible to degradation, such that they were still efficiently degraded by the small amounts of RNase E present in the knockdown condition.

Leadered transcripts appeared to be more sensitive to RNase E levels than leaderless transcripts, suggesting that 5’ UTRs may serve as platforms for engagement with RNase E. However, there was no correlation between degree of stabilization upon *rne* knockdown and predicted secondary structure near the 5’ ends of transcripts. This suggests that the effects of 5’ UTRs on RNase E engagement cannot be explained simply by availability of 5’ ends. This finding is somewhat surprising given the reported strong effect of 5’ end engagement on RNase E activity in *E. coli* (33,49–51), and reports of 5’ end secondary structure protecting transcripts from degradation in *E. coli* (38,52). It is possible that mycobacterial RNase E is less 5’-end dependent than *E. coli* RNase E, or that other transcript features are more important determinants of sensitivity to RNase E.

Our data implicate RNase E as the enzyme responsible for mRNA cleavage events that produce 5’ ends with monophosphorylated cytidines, which are widespread in vivo in both *M. smegmatis* (17) and *M. tuberculosis*. This cleavage sequence preference differs from what was reported in a previous study of the in vitro activity of *M. tuberculosis* RNase E (18). In that study, the presence of a single cytidine in an otherwise mono-uridine oligo was inhibitory to cleavage. However, the effects of cytidines in other sequence contexts were not tested. Our results are therefore not inconsistent with that study, but rather expand upon it. The strong preference of mycobacterial RNase E to cleave 5’ of cytidines contrasts with the lack of strong base specificity by *E. coli* and *Synechocystis* sp. PCC 6803 RNase E at the +1 position ((53) and reviewed in (12)). Residue F67 in *E. coli* RNase E is highly conserved among the Proteobacteria and was proposed to play a key role in the catalytic mechanism by forming a binding pocket for the base one or two nt downstream of the cleavage site (34). Mutating this residue to Ala in *E. coli* abolished activity in vitro (34). However, the residue at the equivalent position in both *M. smegmatis* and *M. tuberculosis* is Val. It is tempting to speculate that differences in the key residues that position the RNA substrate in the active site are responsible for the differences in cleavage sequence preference for mycobacterial vs *E. coli* RNase E. Further work is needed to investigate this question.

Both our in vivo and in vitro data indicate that while RNase E has a strong preference for cleaving 5’ of cytidines, the impact of the surrounding sequence is weak. This could mean that the identities of the surrounding nt are unimportant for RNase E binding and cleavage, or that the identities of those nt are important but act in combinatorial ways that are not obvious from the data currently available. Interpretation of the in vivo cleavage patterns is complicated because (1) cleavage is likely affected by ribosomes and RNA-binding proteins that protect or expose particular regions and (2) cleavage products that are rapidly degraded are not detected and our methods therefore are biased towards identification of cleavage events that produce stable products. In vitro, there was a clear preference for cleavage 5’ of cytidines, but there were many cytidines that did not produce detectable cleavage products, indicating that RNase E prefers certain positions within the test substrate. We examined secondary structure predictions of the substrate and found that the cleaved positions did not correspond to the positions most likely to be in single-stranded loops. The in vitro cleavage pattern therefore cannot be easily explained by the predicted secondary structure. Stem-loops near cleavage sites have been shown to stimulate or direct cleavage by *E. coli* RNase E in some contexts (54–56), and therefore the sites cleaved in our study could be dictated in part by such cis-acting elements. Cis-acting unpaired regions have also been shown to affect cleavage by *E. coli* RNase E (57). The potential impact of the scaffold domains (which were partially deleted in our purified RNase E) should also be considered, as the *E. coli* RNase E scaffold domains were recently shown to affect catalytic activity (58).

Our study highlights the differences in the types of data obtained from different methods of RNA cleavage-site analysis, as well as some of the challenges in identifying RNA cleavage sites. Ligation-based methods, as we used here for *M. tuberculosis* and as we and many others have used in the past for other bacteria, precisely reveal 5’ ends generated by RNA cleavage. However, 5’ ends are only detected from cleavage events that produce relatively stable fragments with sequence and secondary structure characteristics amenable to ligation. Fragments 5’ of cleavage sites are not captured at all; these can be captured by 3- end ligation approaches, but analysis of the resulting datasets is challenging because 3’ ends generated by many RNases (including RNases E, J, and III) are chemically indistinguishable from 3’ ends generated by transcription termination. The ligation-independent method reported previously (37) and modified here, in contrast, does not identify precise cleavage site locations but may give a broader view of the ubiquity and sequence context of cleavage sites attributable to a particular RNase engineered to be induced or repressed. Ligation-based methods may be more useful for identifying cleavage products that are stable and functional, while the ligation-independent approach may provide a more accurate view of the breadth of action of RNases of interest.

It is important to note that for both methods, there is no readily definable cutoff for identifying cleavage sites. It is therefore not possible to conclusively determine the total number of cleavage sites in a transcriptome using the combination of methods we have employed. Using read depth filters similar those we previously published for *M. smegmatis*, here we found ∼3000 high-confidence *M. tuberculosis* cleavage sites with the ligation-based method. Relaxing one of the filters produced a set of ∼10,000 putative cleavage sites with a similar but slightly weaker sequence context signature. Our data suggest a scenario in which the transcriptome contains many cleavage sites, some that are cleaved frequently and/or produce relative long-lived products, and others that are cleaved infrequently and/or produce relatively short-lived products. If this is true, further relaxing the filters would likely reveal still more cleavage sites, likely mixed with a greater proportion of false positives. Some sites may be cleaved so infrequently that their products are not distinguishable from noise. Together, this is consistent with (1) the underlying biology of RNases that have low sequence specificity and/or cleave at ubiquitous sequences (eg, upstream of a cytidine), (2) the fact that mRNA cleavage in vivo is affected by binding of macromolecules such as ribosomes and sRNAs, and (3) the reality that some cleavage products are extremely short-lived and difficult to detect by any method.

It is notable that RNase J, a bifunction endo/exonuclease, did not impact the abundance of most transcript 5’ ends in *M. tuberculosis*. This is consistent with the idea that RNase J has a specialized role in degradation of specific types of highly structured transcripts, as we recently reported (3), rather than a global role. It is also consistent with the idea that RNase J and RNase E may cleave similar sequences (59,60) and therefore have partially redundant activities; however, the start contrast in phenotypes observed in mycobacterial RNase J knockout strains and RNase E knockdown strains suggest such redundancy is limited.

The cleavage sites mapped in *M. tuberculosis* were disproportionately located in untranslated regions. This may reflect the greater accessibility of such regions to RNases, as they lack protection by ribosomes. An intriguing question arising from this observation is the extent to which proteins are produced from translation of cleaved mRNAs. This has been reported in some bacteria, where there are known examples of polycistronic transcripts that are cleaved to produce fragments with different stabilities, leading in some cases to different stoichiometries of proteins encoded in operons (42–48). There is one reported example in mycobacteria but the evidence supporting it are less conclusive (61). Further studies are therefore needed to investigate the functional consequences of stable RNA cleavage products.

## MATERIALS AND METHODS

### Bacterial strains and culture conditions

*Mycolicibacterium smegmatis* strain mc^2^155 and derivatives (Table 1) were grown in Middlebrook 7H9 liquid medium supplemented with glycerol, Tween-80, catalase, glucose, and sodium chloride as described (20) or on Middlebrook 7H10 with the same supplements except for Tween-80. *Mycobacterium tuberculosis* strain H37Rv was grown in the same way with the addition of oleic acid. *Escherichia coli* NEB- 5-alpha (New England Biolabs) was used for cloning and BL21(DE3) pLysS was used for protein overexpression. *E. coli* was grown on LB. Liquid cultures were grown at 37°C with a shaker speed of 200 RPM, except for *M. tuberculosis* which was shaken at 125 RPM. When indicated, anhydrotetracycline was used at 200 ng/mL. Antibiotic concentrations used for mycobacteria were 25 µg/mL kanamycin and 150 µg/mL hygromycin. Antibiotic concentrations used for *E. coli* were 50 µg/mL kanamycin, 150 µg/mL hygromycin, and 34 µg/mL chloramphenicol.

### *M. smegmatis* strain construction

#### SS-M_0418

The repressible *rne* strain was built by mycobacterial recombineering as described (62). A gene replacement cassette was assembled in plasmid pSS187 by NEBuilder HiFi assembly (NEB) and amplified from the plasmid as a linear fragment by PCR. The *rne* TSS is located 236 nt upstream of the translation start site (17), and the core promoter sequence is evident shortly upstream of the TSS as expected. The gene replacement cassette contained nt -846 through -347 relative to the *rne* (msmeg_4626) translation start site (a 500 bp region located upstream of the *rne* native promoter), a hygromycin resistance gene and promoter, the P766(8G) promoter which contains tet operators (tetO), the P766(8G)-associated 5’ UTR, and the first 500 bp of *rne* coding sequence. 2 μg the gene replacement cassette were dialyzed in pure water before transformation into SS-M_0078 (WT *M. smegmatis* with the recombinase plasmid pNit-recET-Kan). Correct integration of this cassette replaced the 346 nt upstream of the *rne* translation start site with the hyg resistance gene and the P766(8G) promoter and 5’ UTR, and was confirmed by sequencing. Counterselection with 15% sucrose was followed by PCR screening to identify an isolate (SS-M_0151) that lost the recombinase plasmid. SS-M_0151 was further transformed with plasmid pSS291 encoding a Tet repressor (TetR) into the L5 phage integration site.

#### SS-M_0424

A hygromycin-resistant control strain was built using the method described for SS-M_0418, the difference being that the target DNA fragment that was transformed into SS-M_0078 only contained the hygromycin resistance cassette with sequence upstream and downstream of position -346 relative to the *rne* translation start site, resulting in insertion of the hyg resistance gene and promoter without deletion of any native sequence.

### RNA extraction, RNAseq library construction, and sequencing

Cultures were grown to an OD of 0.8-0.9, with or without addition of ATc 8 hrs prior, and divided into a series of 14 ml conical tubes. RIF was added to a final concentration of 150 µg/mL and cultures were harvested after 0, 1, 2, 4, 8, 16, or 32 min by freezing in liquid nitrogen. Frozen cultures were stored at - 80°C and thawed on ice for RNA extraction. RNA was extracted as in (20). Illumina libraries were constructed and sequenced by the Broad Institute Microbial ‘Omics Core using the library construction procedure described in (63).

### cDNA synthesis and quantitative PCR

cDNA was synthesized as described (20) and qPCR was performed using the conditions described in (20) and the primers listed in that work and in Table S6.

### Gene re-annotations in M. smegmatis and M. tuberculosis

For *Mycolicibacterium smegmatis*, we used the genome sequence of *M. smegmatis* mc^2^155 strain (NC_008596.1) from Mycobrowser Release 4 (64). For gene annotations, we combined all the annotations from PATRIC 3.6.10 (65), Mycobrowser Release 4 (64) and recently identified novel ORFs (17). The combined annotations were first updated with reannotations of 213 genes as previously described (17). Based on the assumption that transcripts starting with AUG or GUG will be translated in a leaderless fashion (40), we then further utilized the transcriptional start sites (TSS) reported in (17) to re-annotate 156 genes whose annotated 5’ UTRs started with in-frame AUG or GUG codons. In these cases the coding sequence was re-annotated to start at the TSS. The resulting annotations were scrutinized to exclude duplications and genes with frame shift errors. The reannotated CDS boundaries are listed in Table S8 and were used for all further analyses unless stated otherwise.

For *Mycobacterium tuberculosis*, the genome sequence and original gene annotations of *M. tuberculosis* H37Rv strain (NC_000962.3) were obtained from Mycobrowser Release 4 (64). Then for genes with only one defined TSS, we used the following procedure to determine if the coding sequence starting coordinates would be re-annotated (40). For genes with TSSs upstream of the previously annotated start codon, we re-annotated the start of the coding sequence to the TSS for those genes with in-frame AUG or GUG at the 5’ end of the transcript. For genes that had a single TSS downstream of the previously annotated start codon, the start of the coding sequence was re-annotated to the position of the TSS if the TSS was at an in-frame AUG or GUG within the first 30% of the previously annotated coding sequence. If the TSS was not at an in-frame AUG or GUG, we re-annotated the start of the coding sequence only if the next in-frame start codon (AUG, GUG, or UUG) was found in the first 30% of the previously annotated coding sequence. The reannotated CDS boundaries are listed in Table S9 and were used for all further analyses unless stated otherwise.

### RNAseq data analysis for differential expression analysis

The 0-min RIF-treated samples were used to measure and compare steady-state transcript abundance. Reads were aligned to *M. smegmatis* mc^2^155 reference sequence NC_008596.1 from Mycobrowser Release 4 (64) with Bowtie v1.2.2 (66), read alignment processed by SAMtools v1.9 (67), counts determined by HTSeq v0.10.1 (68). The differential expression analysis was performed using Clipper with the gene counts normalized by qPCR normalization factors (35).

### Gene set enrichment analysis

The enrichment of KEGG pathway was tested using ClusterProfiler v4.4.4 (36), based on the gene list sorted by log_2_ fold changes of expected and observed abundance. FDR-adjusted *p*-values were used for multiple testing correction.

### RNAseq data analysis for expression-library-based cleavage site analysis in *M. smegmatis* and *M. tuberculosis*

This analysis was performed on the *M. smegmatis* 0-min RIF-treated samples as well as an *M. tuberculosis rne* knock down strain and corresponding control strain ((16), GEO accession GSE126286). Quality control was performed using FastQC. Reads were first scanned from 5’ end to 3’ end and cut once the average quality per base of 4-base wide sliding window dropped below 20. After such processing, reads with less than 25 bases were discarded using Trimmomatic v0.39 (69). Reads were aligned using Bowtie2 v2.4.5 (70) with the “--very-sensitive” option. We first aligned reads to tRNA and rRNA sequences only. The remaining reads were aligned to NC_008596.1 (*M. smegmatis*) or NC_000962.3 (*M. tuberculosis*). Via SAMtools v1.16.1 (67), we filtered the resulting alignments by keeping only the primary alignments with MAPQ at least 10. The aligned, filtered reads that mapped in proper pairs were split into their corresponding strands to quantify strand-specific coverage at the single-nucleotide level using BEDTools v2.30.0 (71). The coverage for each gene was then calculated by summing the single-nucleotide coordinate coverage within the gene, and the average nucleotide coverage for each gene was calculated by dividing the summed coverage by gene length. We only kept genes with average nucleotide coverage at least 5 in all replicates and conditions. For those qualified genes, we excluded nucleotides at overlapped gene regions for downstream analysis. To correct for the variability in expression level among genes and between conditions, we normalized single-nucleotide coverages using the whole-gene coverages. The single-nucleotide coverages were divided by the total summed nucleotide coverage of each gene (excluding regions overlapping other genes) after adding one pseudocount to all nucleotide positions. The final normalized coverage of each nucleotide was the average of triplicates in each condition.

The coverage ratio at each qualified nucleotide position between any two conditions was then calculated as the log_2_(Condition1/Condition2) ratio. For each nucleotide under investigation, we quantified the sequence context using the relative nucleotide frequency of the 20 bases upstream and downstream.

### RNAseq data analysis for determination of half-lives in *M. smegmatis*

To calculate mRNA half-lives, data from all of the timepoints following RIF treatment were processed. First, reads were aligned using BWA-MEM v0.7.17 (Li, 2013). Next, the resulting alignments were processed for each strand by SAMtools v1.10 (Danecek et al., 2021). The raw coverage of each coordinate was calculated through BEDTools v2.29.1 (Quinlan & Hall, 2010). Then we conducted a two-step normalization of the raw coverage. First, coverage was normalized by the total number of reads in each library. Then we calculated normalization factors by performing qPCR to determine the relative expression levels of eight genes (*sigA*, *rraA*, *esxB*, *atpE*, *rne*, msmeg_4665, msmeg_5691, msmeg_6941; Table S7) at each sample and timepoint compared to the average of the 0-min RIF control strain (no ATc) samples. qPCR was done with cDNA made from random priming as described above, separately from RNAseq library construction. Each qPCR reaction was performed using 400 pg of cDNA. As ribosomal rRNA depletion was not performed, the CTs obtained from the qPCR reflect the expression level of the target gene relative to the total RNA pool, which is primarily rRNA. Normalization factors were calculated separately for the region amplified by qPCR in each of the eight genes and averaged. Specifically, for a given sample *T_n_*, we calculated the normalization factor *F_Tn_* from the qPCR target gene expression measurements as indicated below:

Calculation of the expected RNAseq coverage (*T_n,i,RNAseq_expected_*) for each qPCR amplicon region (*_i_*) in each sample (*T_n_*), where *T_0_* represents the average value for the control strain without ATc immediately after addition of RIF, and *qPCR* represents relative abundance of the amplicon determined by qPCR:

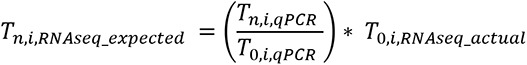

Calculation of a global normalization factor (*F_Tn_*) by calculating and averaging the normalization factors for each qPCR amplicon region:

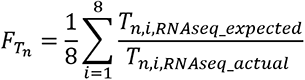

Then the final normalized coverage for each coordinate was calculated by multiplying the first step normalized coverage by the global normalization factor for each sample. The coverage for each gene was then represented by the summation of the normalized coverage of its coordinates, divided by the gene length.

### Calculation of predicted transcription rates

Predicted transcription rates were calculated as a function of steady-state abundance and mRNA degradation rate as described (72) and as follows:

Transcription rate = VT = (k*mRNA) + (μ*mRNA)
mRNA = steady-state mRNA abundance (taken from 0 min RIF treatment)
k = degradation rate = ln(2)/half life
μ = growth rate = ln(2)/doubling time
Doubling time = 150 min

The predicted transcription rate units are arbitrary and therefore useful only for comparison of genes or conditions.

### mRNA cleavage site mapping in *M. tuberculosis*

Mapping of *M. tuberculosis* TSSs was previously described (40). The same dataset was used to identify mRNA cleavage sites. All analyses of this dataset were done using the genome annotations in NC_000962.gbk rather than the reannotations shown in Table 9. As described in (40), RNA 5’ ends were identified, filtered based on absolute read depth and read depth relative to local expression library coverage, and subject to gaussian mixture modeling to distinguish between TSSs and cleavage sites on the basis of relative coverage in libraries from RNA treated with RppH (“converted,” capturing both TSSs and cleavage sites) and libraries from untreated RNA (“non-converted,” capturing primarily cleavage sites). 5’ ends with converted/non-converted library read depth ratios less than 1.39 had a cumulative probability of ≤0.01 of belonging to the TSS population (after adjusting for multiple comparisons by the Benjamini-Hochberg procedure) and were therefore designated RNA cleavage sites. Because cleavage may be imprecise, filtering was performed to retain the single cleavage site with the greatest converted-library read coverage in each 5 nt window. This resulted in the 2983 high-confidence cleavage sites reported in Table S5A. The longer list of putative cleavage sites reported in Table S5I was obtained by applying the same converted/non-converted ratio cutoff to a list of 5’ ends from earlier in the pipeline prior to filtering on coverage relative to local expression library coverage. Instead, only a filter requiring a minimum mean converted library read depth of 20 was applied. This resulted in 10795 putative cleavage sites.

### M. tuberculosis TSS analyses

TSSs from the above dataset were considered to be associated with the 5’ ends of genes if they were either (1) within 500 nt upstream of an annotated start codon or (2) within the first 25% of an annotated coding sequence. TSSs were considered to be internal within coding sequences if they were located between 25% and 80% of the way through annotated coding sequences. TSSs were considered to be associated with putative antisense transcripts if they did not meet any of the above criteria and were either (1) located on the opposite strand of an annotated coding sequence or (2) located <200 nt from the end of an annotated coding sequence on the opposite strand. TSSs were considered to be intergenic if they did not meet any of the above criteria for 5’-end associated, internal, or antisense transcripts.

Genes were assigned to operons if they were transcribed consecutively on the same strand and if both of the following criteria were met: (1) Only the first gene had an assigned TSS and (2) The downstream gene(s) were sufficiently expressed. Sufficient expression was defined as having a Reads Per Kilobase of transcript per Million mapped reads (RPKM) value in corresponding RNA-seq expression libraries equal to the 5th percentile or above of RPKM values for all genes with TSSs. This prediction algorithm is conservative and excludes many loci that may be transcribed both polycistronically and monocistronically.

### Secondary structure prediction

Free energy of RNA folding and basepair probabilities for minimum free energy structure were predicted using the Vienna RNA Package utility RNAfold (73). For Figure 6B, the 200 nt region spanning each RNA cleavage site was extracted and the minimum free energy of secondary structure formation was predicted for 39 nt sliding windows across each such region. The data plotted are the mean and the 25th and 75th percentile minimum free energies of 39 nt windows centered around each relative coordinate in all cleaved RNAs.

### 5’ RACE to map a putative RNase E cleavage site in the rRNA transcript

Enzymes were obtained from New England Biolabs unless otherwise specified. Five hundred ng of each RNA sample were mixed with 1 µg of oligo SSS1016 in a total volume of 9 µl, incubated at 65°C for 10 minutes, and cooled on ice for 5 minutes. Each sample was combined with 21 µl of ligation mix containing 10 µl of 50% PEG8000, 3 µl of 10X T4 RNA ligase buffer, 3 µl of 10 mM ATP, 3 µl of DMSO, 1 µl of murine RNase inhibitor, and 1 µl of T4 RNA ligase. Samples were incubated at 20°C overnight and purified with a Zymo RNA Clean & Concentrator-5 kit according to the manufacturer’s instructions with the following modifications: samples were first diluted by addition of 20 µl of RNase-free water, and samples were eluted in 8 µl of RNase-free water. Three µl of each purified ligation were then subject to cDNA synthesis or mock (no-RT) cDNA synthesis. Samples were combined with 1 µl of a mix containing 50 mM Tris pH 7.5 and 500 ng/µl random primers (Invitrogen), incubated at 70°C for 10 minutes, and snap-cooled in an ice-water bath. cDNA synthesis was done as described (20). 35 ng of cDNA or the equivalent volume of the corresponding no-RT sample were mixed with 2.5 µl 10X Taq buffer, 1.25 µl each 10 µM primers SSS1017 and SSS2210, 1.25 µl DMSO, 0.5 µl of 10 mM each dNTP mix, 0.167 µl Taq polymerase, and water to a final volume of 25 µl. Cycling conditions were 5 minutes at 95°C, 35 cycles of 30 seconds at 95°C, 20 seconds at 52°C, and 25 seconds at 68°C, and a final 5 minute incubation at 68°C. PCRs were run on 1.5% agarose gels and bands that appeared in cDNA samples but not in no-RT samples were excised and sequenced with SSS2210 to identify the adapter/RNA junctions.

### Overexpression and purification of recombinant RNase E variants

Two RNase E variants were recombinantly expressed and purified for in vitro RNA cleavage assays: residues 146-824 (partial N-terminal truncation and full C-terminal truncation), and residues 146-824 with D694R and D738R mutations. pSS348, carrying the *M. smegmatis rne* coding sequence with a Δ1-145aa partial N-terminal deletion, Δ825-1037aa full C-terminal deletion, and an N-terminal addition of 6XHis tag, 3XFLAG tag, TEV protease cleavage site, and 4XGly linker sequences, was used as a template for creation of pSS420, which encodes RNase E residues 146-824 with the indicated tags in a pET38 backbone. pSS420 was then used as a template for creation of pSS421, which has the mutations D694R and D738R, predicted to abolish catalytic activity (34). All constructs were sequenced to confirm the success of point mutations and truncations.

*E. coli* strain BL21(DE3)pLysS was transformed with each of the RNase E expression plasmids and 500 mL cultures were grown to an OD600 of ∼0.5, then induced with 400 *μ*M IPTG and incubated at 28°C for four hours prior to harvest. Pellets were resuspended in 1X IMAC buffer (20 mM Tris-HCl pH 7.9, 150 mM NaCl, 5% glycerol, 0.01% Igepal) containing 10mM imidazole and lysed with a BioSpec Tissue-Tearor (10 cycles of 15 seconds each at maximum speed, with 60 seconds on ice between cycles). Lysates were cleared by centrifugation, incubated for 30 minutes on ice with His-Pur Ni-NTA resin (Thermo Scientific), packed to a height of approximately 5 cm in polypropylene columns, washed with IMAC buffer containing 10 mM imidazole, and eluted with IMAC buffer containing 150 mM imidazole. Eluates were concentrated with Microcon PL-30 (30,000 NMWL) protein concentrators (Millipore Sigma) and loaded onto 1 cm diameter, 38 mL Sephacryl S-200 High Resolution resin (GE Healthcare) size exclusion chromatography columns. Flow rate was regulated using a Masterflex C/L pump. The buffer was 1X IMAC with the addition of 1 mM EDTA and 1 mM DTT.

### Preparation of in vitro-transcribed RNA substrates

Genomic DNA was used as a template to produce PCR products containing portions of the *atpB-atpE* locus downstream of the T7 Phi2.5 promoter and sequence needed for A-initiated transcription (TAATACGACTCACTATT**A**GG, where transcription initiates at the bolded “A”). One PCR product had the promoter oriented to produce the sense strand, and the other was shorter and had the promoter oriented to produce a partial antisense strand (Fig. S12). Monophosphorylated RNA was synthesized from each of these PCR products in the presence of a 50-fold molar excess of AMP over ATP (74) with T7 RNA polymerase (NEB M0251). Each 50 μL reaction contained 1X reaction buffer, 5 mM DTT, 1 mM UTP, 1 mM CTP, 1 mM GTP, 0.5 mM ATP, 25 mM AMP, 5 units/μL T7 RNA polymerase, 1 unit/μL Murine RNase inhibitor, and 2 μg DNA template. Reactions were incubated at 37°C for 16 hours. The resulting transcripts were treated with TURBO DNase at 37°C for 30 minutes before purification with a Zymo RNA Clean & Concentrator-5 kit.

The *atpB-E* sense transcript and anti-sense transcript were combined at a 1:1 molar ratio and the mixtures were incubated in the presence of 5X annealing buffer (50 mM Tris-HCl, pH 7.9, 0.5 mM EDTA, pH 8.0, 100 mM NaCl) in a 10 μL reaction for 1 min at 90°C, then slowly cooled down to room temperature over a period of approximately 30 min. The resulting annealed RNA mix was immediately stored at -80°C.

### In vitro RNase E cleavage reactions

In vitro RNase E cleavage reactions were heated at 65°C for 3 min prior to adding the enzyme, then cooled and incubated at 37°C for 1 hour following addition of the enzyme. The reaction buffer was composed of 20 mM Tris-HCl, pH 7.9, 100 mM NaCl, 5% Glycerol, 0.01% IGEPAL, 0.1 mM DTT, 10 mM MgCl_2_, and each reaction containing 300 ng annealed RNA mix and 80 ng of purified RNase E. For mock reactions, water was used instead of enzyme. Reactions were stopped by adding equal volumes of 2X Invitrogen^TM^ Gel loading buffer II and then subjected to electrophoresis on a 7.5% or 5% polyacrylamide-8 M urea gels and visualized after 15 min staining with SYBR Gold Nucleic Acid gel stain. Bands of interest were excised, and RNA was recovered using Zymo small-RNA PAGE recovery kit, followed by identification of cleavage sites by 5’ RACE or 3’ RACE.

### 5’ RACE and 3’ RACE to map cleavage sites from in vitro RNase E cleavage reactions

For 5’ RACE, RNA extracted from bands as described above was mixed with 1 μg of RNA oligo SSS1016 in a total volume of 9 μL at 65°C for 5 min, chilled on ice and then combined with 30 U T4 RNA Ligase 1 (NEB M0437M), 40 U Murine RNase Inhibitor (NEB), 10% DMSO, 1 mM ATP, 1X T4 RNase Ligase 1 reaction buffer, and 16.7% PEG 8000 in reactions with a total volume of 30 μL. Reactions were incubated at 20°C for 18 hours followed by column purification. cDNA was synthesized using the reverse oligo SSS916 which anneals close to 3’ end of the sense strand and the cDNA synthesis protocol described above. cDNA was purified and then was used as template to perform Taq PCR with primers SSS1018 and SSS916. Purified PCR products were sequenced with oligo SSS916.

For 3’ RACE, RNA extracted from bands as described above was mixed with 1 μg RNA oligo SSS2433 (which has a 5’ monophosphate and a 3’ inverted deoxythymidine and was modified from (75) at 65°C for 5 min, chilled on ice and incubated at 17°C for 18 hours with the same reaction mix as used for 5’ RACE above. Following column purification, cDNA was synthesized using reverse oligo SSS2434 which anneals to the 3’ adapter, and the protocol described above. cDNA was purified and then was used as template to perform Taq PCR with primers SSS917 and SSS2434. Purified PCR products were sequenced with oligo SSS917.

### Statistical analyses and scripts

Statistics shown in figures 1, 2, and 6 were done in Graphpad Prism version 9.2.0. The scripts for RNAseq processing, analysis and result visualization are available on Github (https://github.com/ssshell/Mycobacterial_RNase_E).

## ACKNOWLEDGMENTS

We thank members of the Shell and Fortune labs for helpful discussions.

## DATA AVAILABILITY

All RNAseq data generated in this study are available at GSE227248. The scripts for RNAseq processing, analysis and result visualization are available on Github (https://github.com/ssshell/Mycobacterial_RNase_E).

## SUPPORTING INFORMATION

Table S1. Half-lives of *M. smegmatis* transcripts with and without *rne* knockdown.

Table S2. Analysis of changes in mRNA half-life and abundance upon *rne* knockdown in *M. smegmatis*.

Table S3. Statistical analysis of *rne* knockdown on mRNA abundance in *M. smegmatis*.

Table S4. Effects of *rne* knockdown on predicted transcription rates in *M. smegmatis*.

Table S5. *M. tuberculosis* cleavage sites and TSS data.

Table S6. Differentially abundant *M. tuberculosis* gene pairs bisected by cleavage sites.

Table S7. Oligos used in this work.

Table S8. *M. smegmatis* gene annotations used in this work.

Table S9. *M. tuberculosis* gene annotations used in this work.

## Supplemental figures

**Figure S1.**
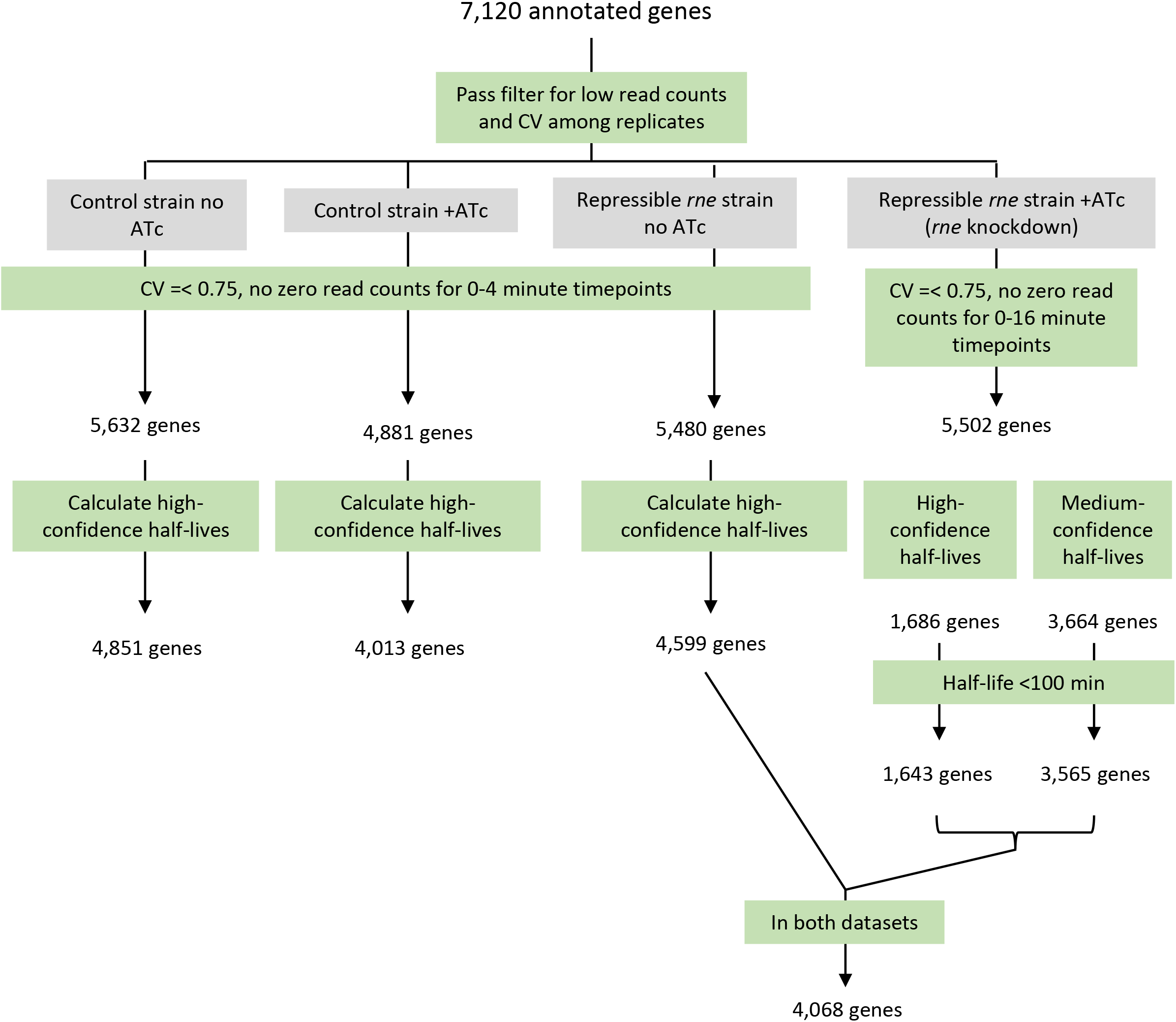
Overview of RNAseq data filtering for half-life calculations. Genes were used when they passed filters for read depth and CV among replicates. Half-life calculations are diagrammed in figures S2 and S3.

**Figure S2.**
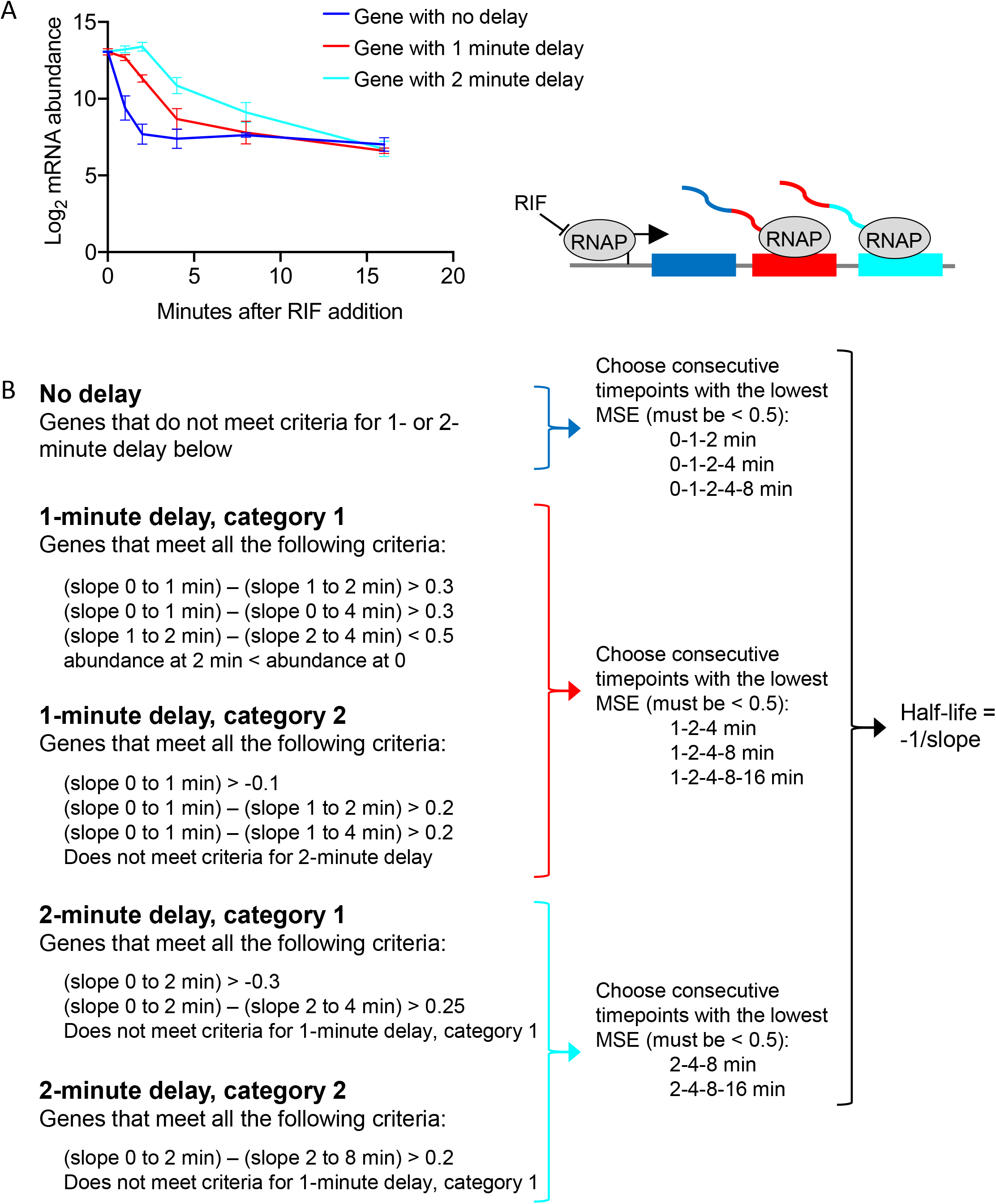
Half-life calculation procedure for genes in control conditions (*rne* not repressed). **A.** Log_2_-transformed mRNA abundance data show three distinct degradation patterns following addition of rifampicin to block transcription. Because rifampicin blocks transcription initiation but not transcription elongation, some genes show a delay before transcript levels decrease. The delay generally corresponds to the distance between the gene and its transcription start site. Furthermore, degradation for all genes reaches a plateau at later timepoints. We expect that the linear portion of the degradation curve between the delay (if present) and plateau is most likely to reflect the true degradation rate and therefore use this to calculate the half-life. **B.** Classification of genes into three delay categories based on linear regression fits to different sets of timepoints following addition of rifampicin, followed by half-life determination. The slopes between the indicated timepoints were calculated and used to classify genes as having no delay, a 1-minute delay, or a 2-minute delay. After removal of early timepoints as indicated to account for the delay, the MSE was used to quantify goodness of fit for linear regression using subsets of the remaining timepoints and the set of timepoints with the best fit were used to calculate the half-life.

**Figure S3.**
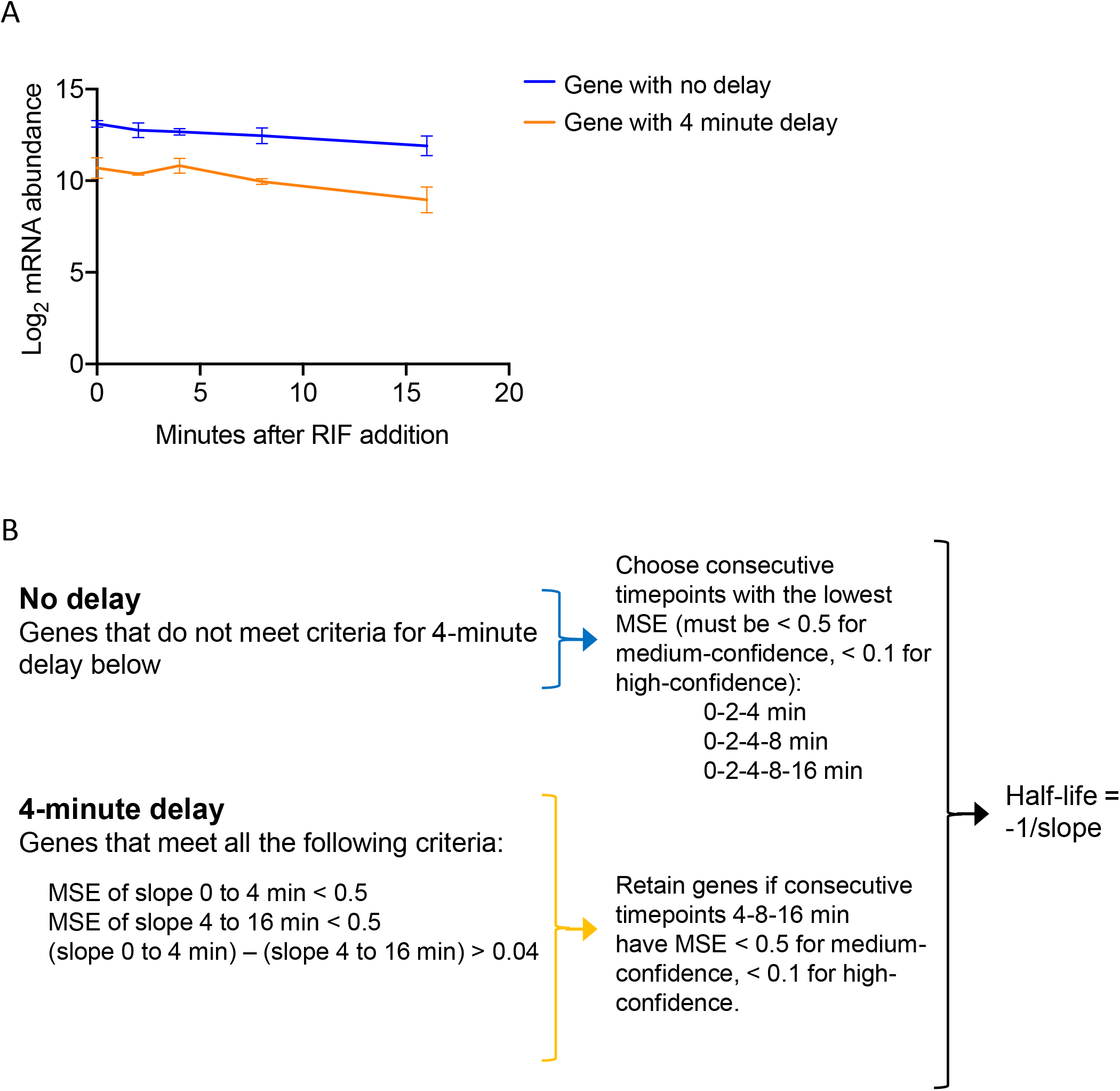
Half-life calculation procedure for genes in *rne* repression condition. **A.** Log_2_-transformed mRNA abundance data show two distinct degradation patterns following addition of rifampicin to block transcription, which can be best categorized as no delay or a 4-minute delay. **B.** Classification of genes into two delay categories based on linear regression fits to different sets of timepoints following addition of rifampicin, followed by half-life determination. The slopes between the indicated timepoints were calculated and used to classify genes as having no delay, or a 4-minute delay. After removal of early timepoints as indicated to account for the delay, the MSE was used to quantify goodness of fit for linear regression using subsets of the remaining timepoints and the set of timepoints with the best fit were used to calculate the half-life.

**Figure S4.**
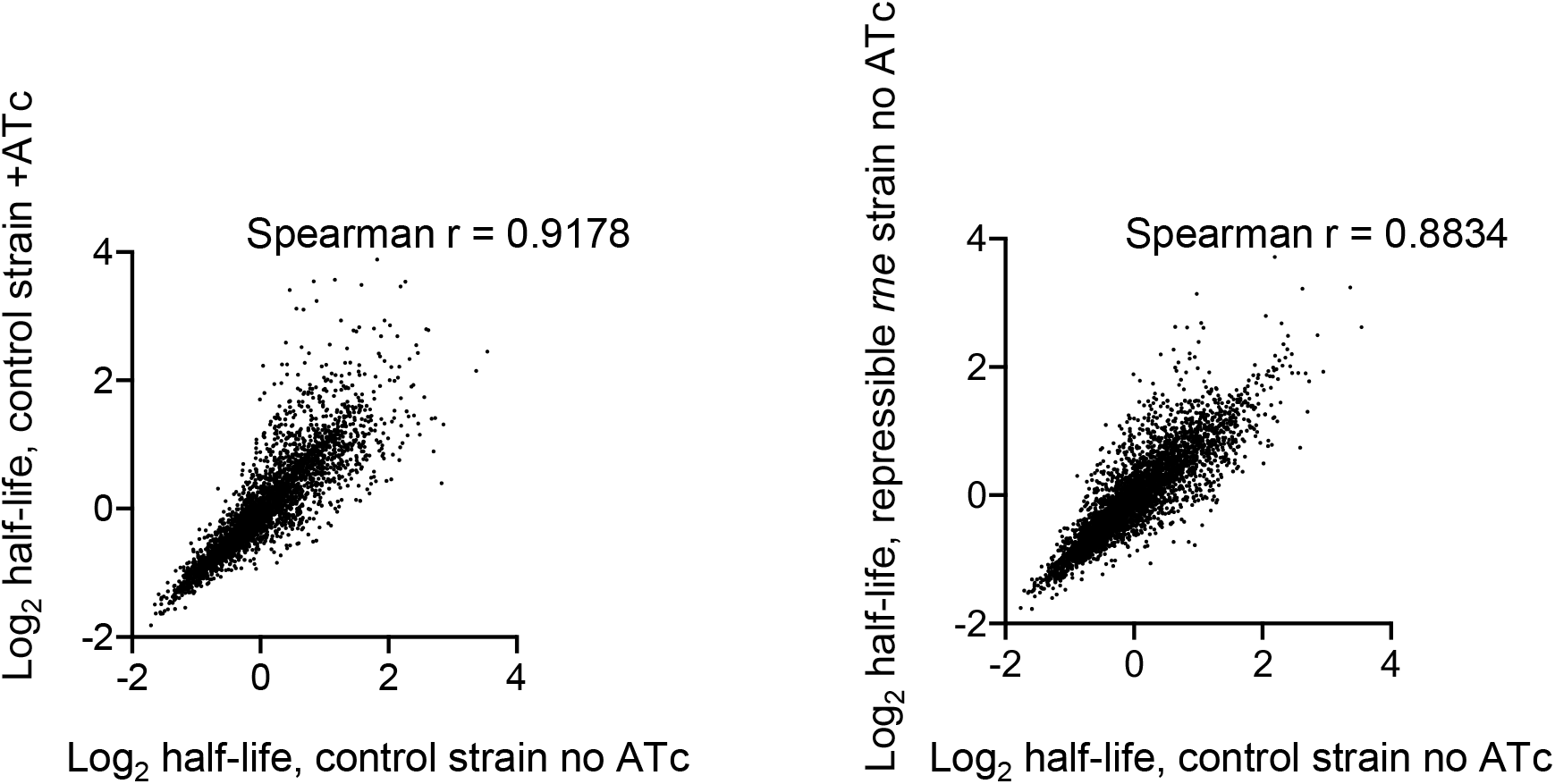
Correlations of half-lives between control conditions. Scatterplots show the half-lives calculated for genes in the control strain with and without ATc (left) and for the repressible and control strains without ATc (right).

**Figure S5.**
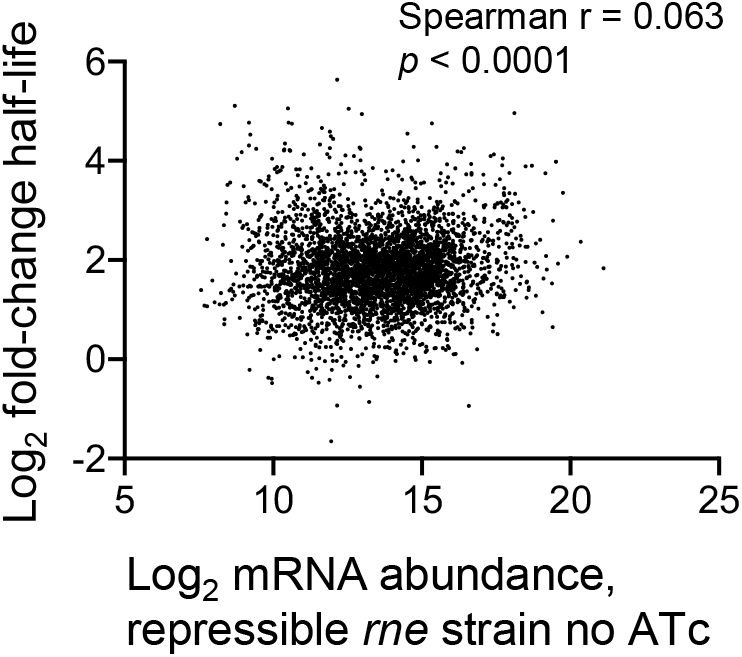
Fold-increase in half-life upon *rne* repression is weakly correlated with abundance prior to repression. Each dot represents a gene for which half-lives were determined in the presence and absence of ATc in the repressible *rne* strain.

**Figure S6.**
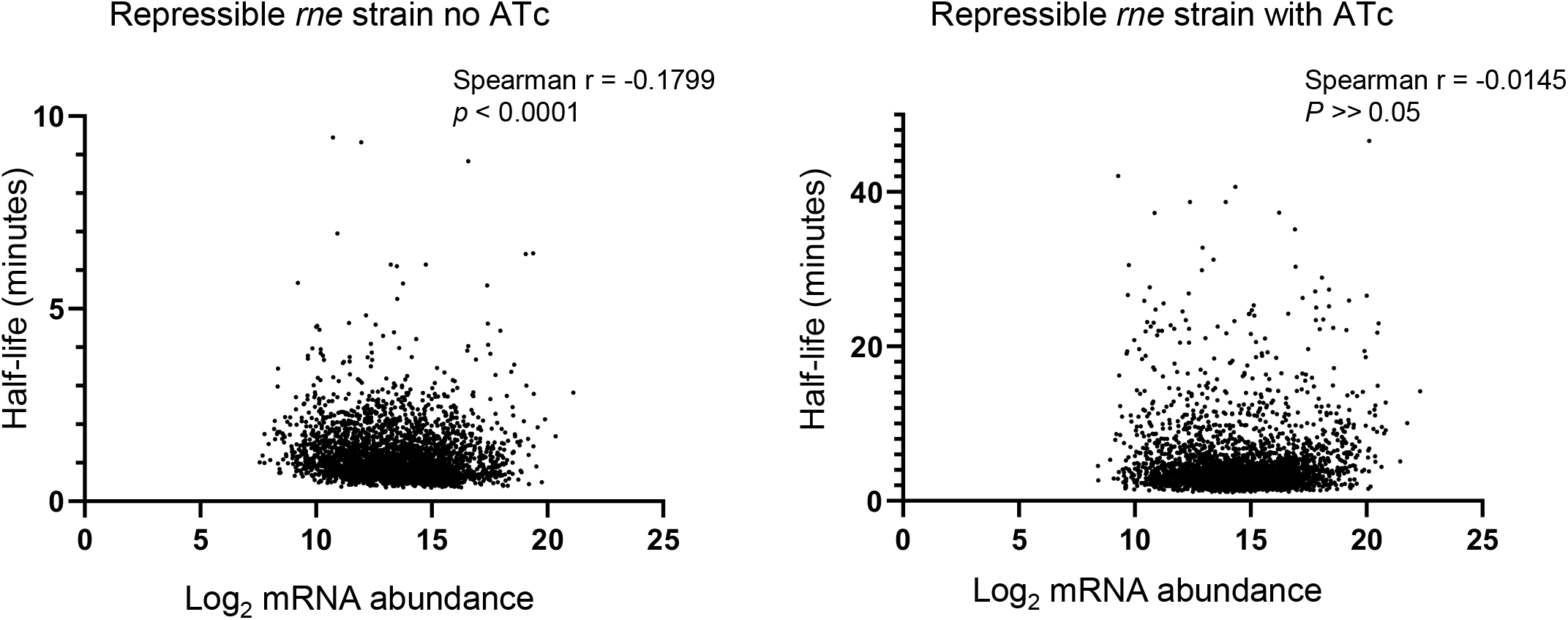
The relationship between mRNA abundance and mRNA half-life changes upon *rne* knockdown. Each dot represents a gene for which half-lives were determined in both the presence and absence of ATc in the repressible *rne* strain.

**Figure S7.**
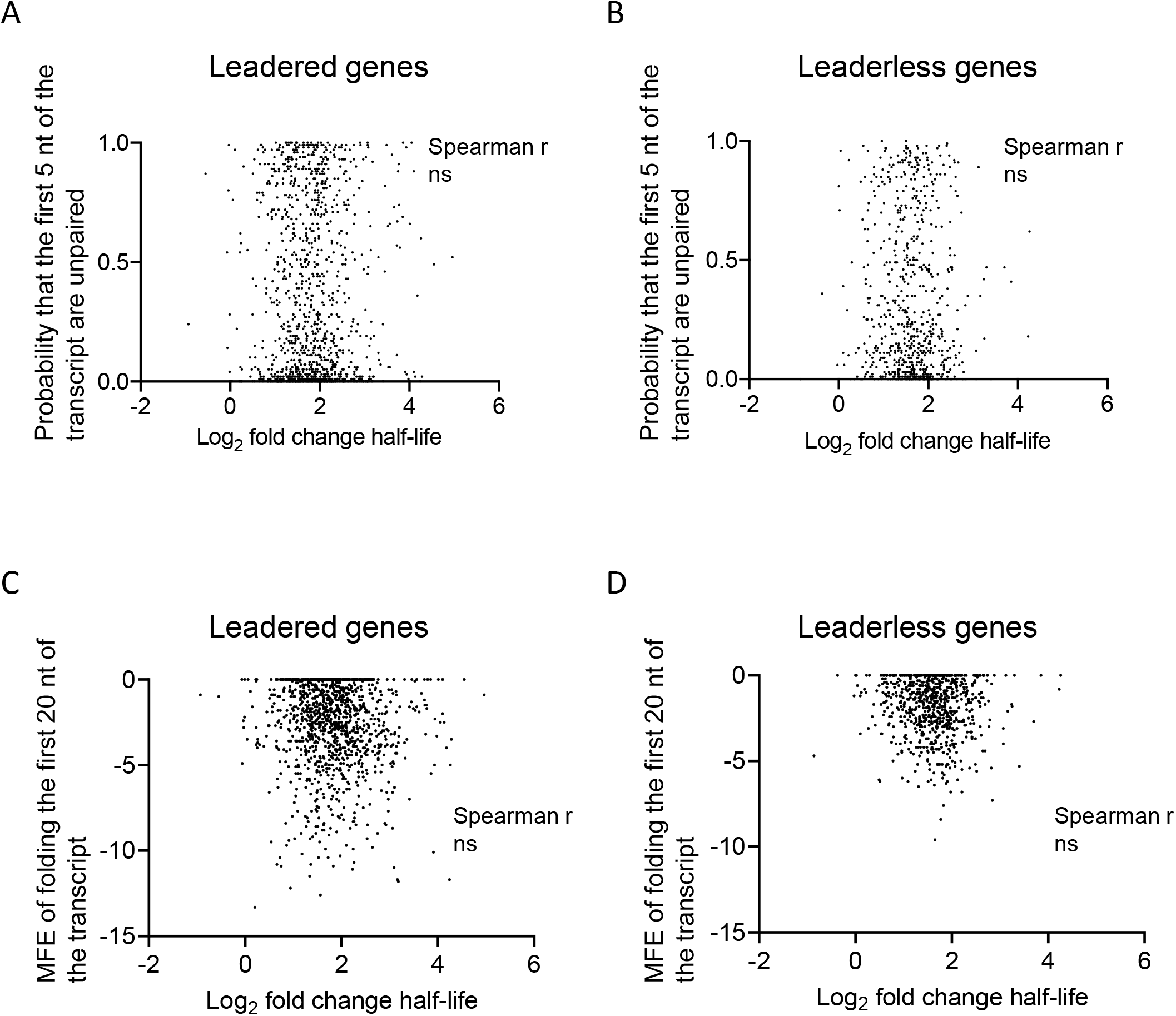
Predicted secondary structure near transcript 5’ ends is not correlated with degree of stabilization upon *rne* repression. Each dot represents a gene for which half-lives were determined in the presence and absence of ATc in the repressible *rne* strain. The MFE structure was predicted for the first 20 nt of each transcript (the 5’ 20 nt of the 5’ UTR for leadered transcripts, and the first 20 nt of the coding sequence for leaderless transcripts). **A** and **B**, the probabilities of the first 5 nt of the transcript being unpaired given the given predicted MFE structures were determined. **C** and **D**, the MFE of folding was determined. All analyses were done in the Vienna RNAfold package. ns, *P* > 0.05.

**Figure S8.**
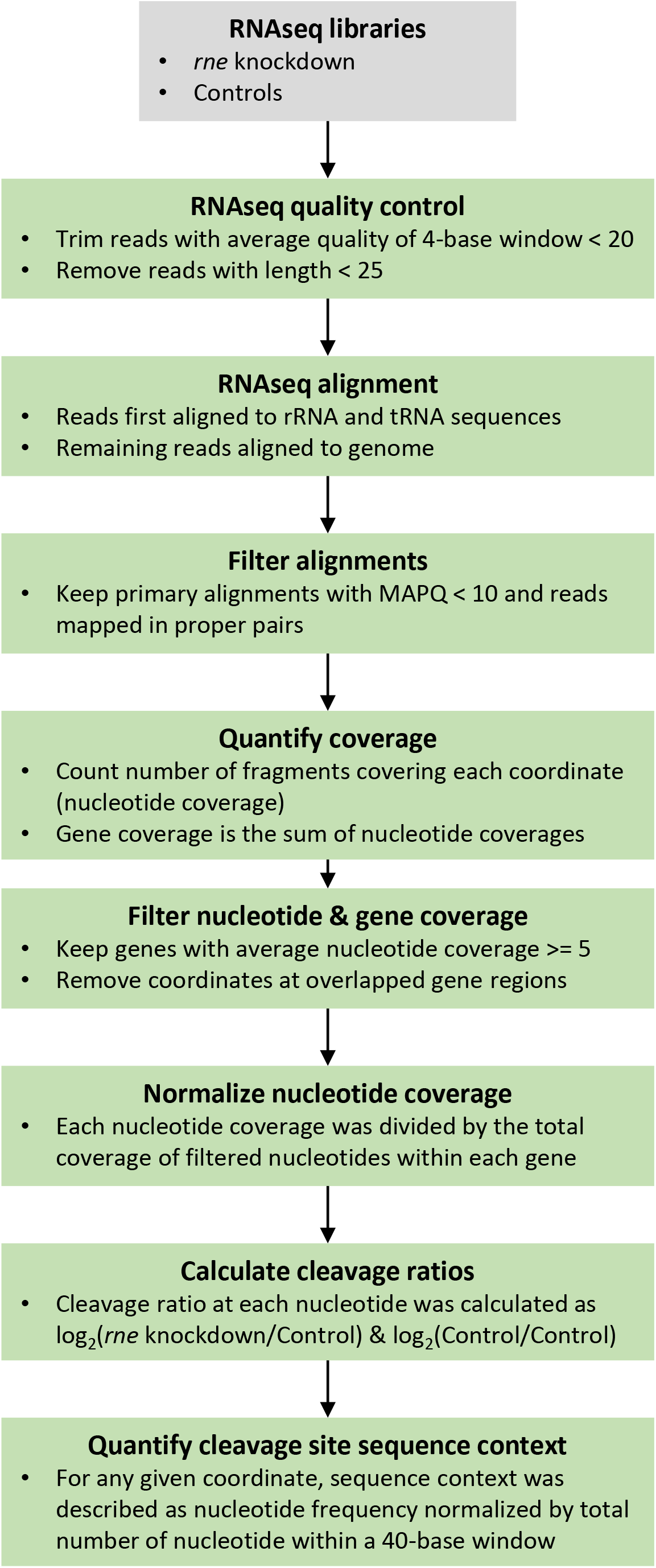
**Pipeline for identifying RNase E cleavage sites from standard Illumina RNAseq expression libraries.**

**Figure S9.**
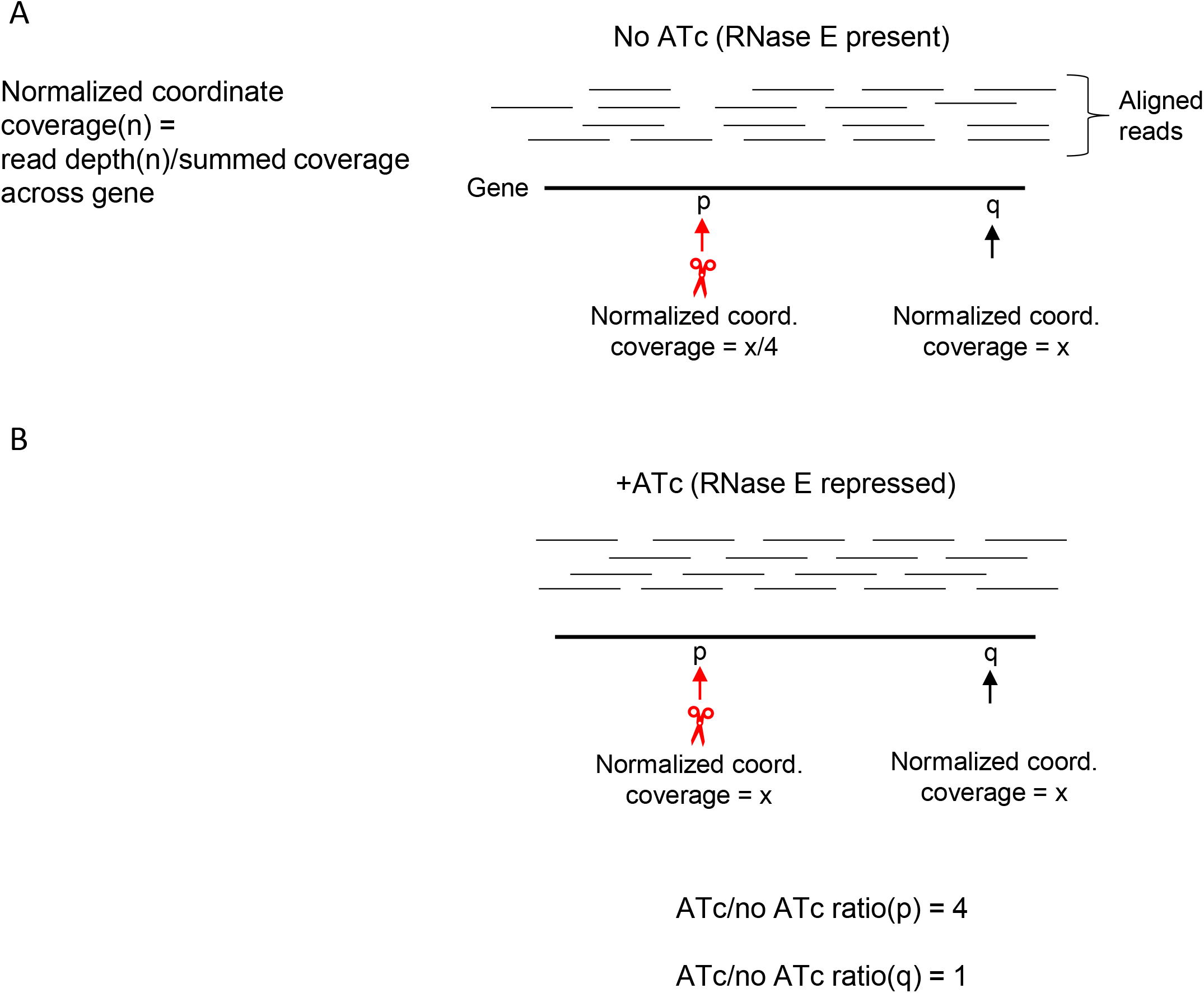
Schematic example of the comparison of read depth in *rne* repressed and non-repressed conditions to identify coordinates whose coverage is affected by RNase E levels. **A.** Within each gene, RNAseq read depth (coverage) at each coordinate is normalized by the summed read count for the entire gene. Coordinate p (at or near a cleavage site) has a lower normalized read depth value than coordinate q (not a cleavage site). **B.** Upon *rne* knockdown, reduced cleavage results in more reads that span the cleavage site. Normalized coverage ratios in the presence vs. absence of ATc are expected to be higher for coordinates at or near RNase E cleavage sites than for coordinates not near cleavage sites.

**Figure S10.**
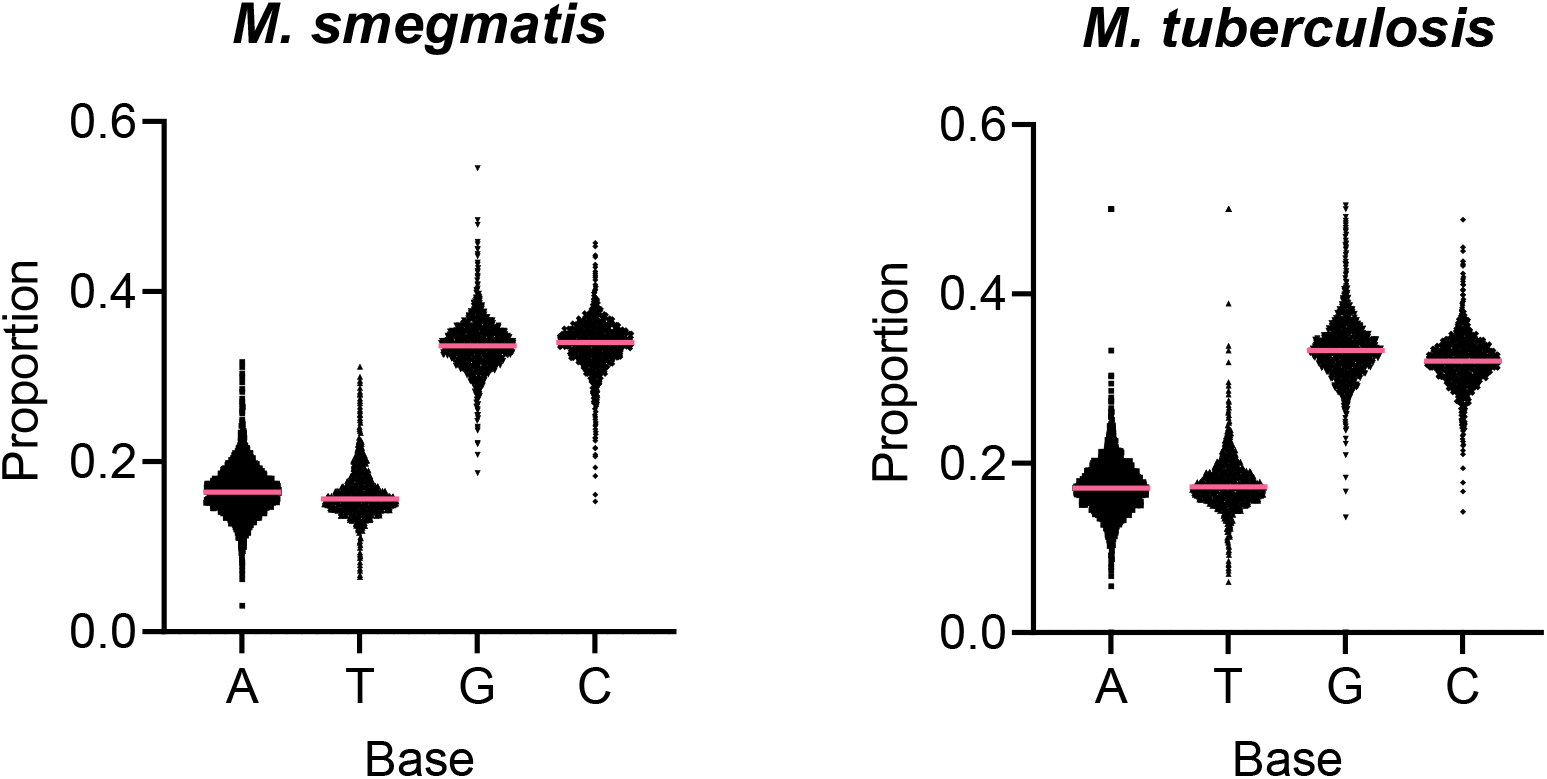
Base composition of coding sequences in *M. smegmatis* and *M. tuberculosis*. Each dot represents a gene. Pink lines indicate medians.

**Figure S11.**
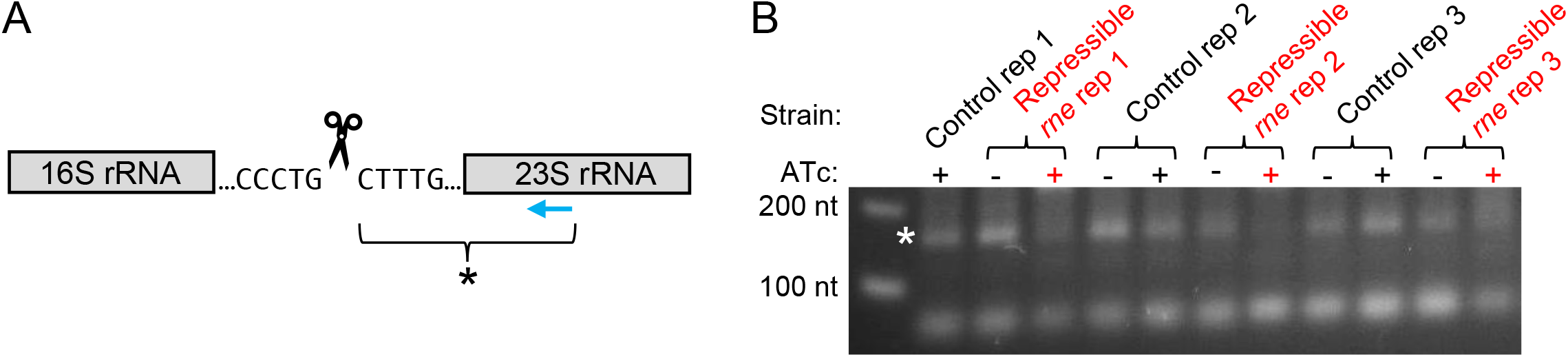
RNase E cleaves upstream of a cytidine during rRNA processing. **A.** Schematic of part of the rRNA operon with the sequence of a region reported to be cleaved by RNase E shown (Taverniti et al 2009). Graphic not to scale. The scissors indicate the exact cleavage position that we mapped by 5’ RACE. The asterisk indicates the 5’ RACE PCR product shown in panel B. The blue arrow indicates the primer used for cDNA synthesis. **B.** An ethidium-bromide strained agarose gel revealing 5’ RACE PCR products. The asterisk indicates the PCR product shown schematically in panel A. Triplicate samples are indicated. Control strain replicate 1 in the absence of ATc was not run.

**Figure S12.**
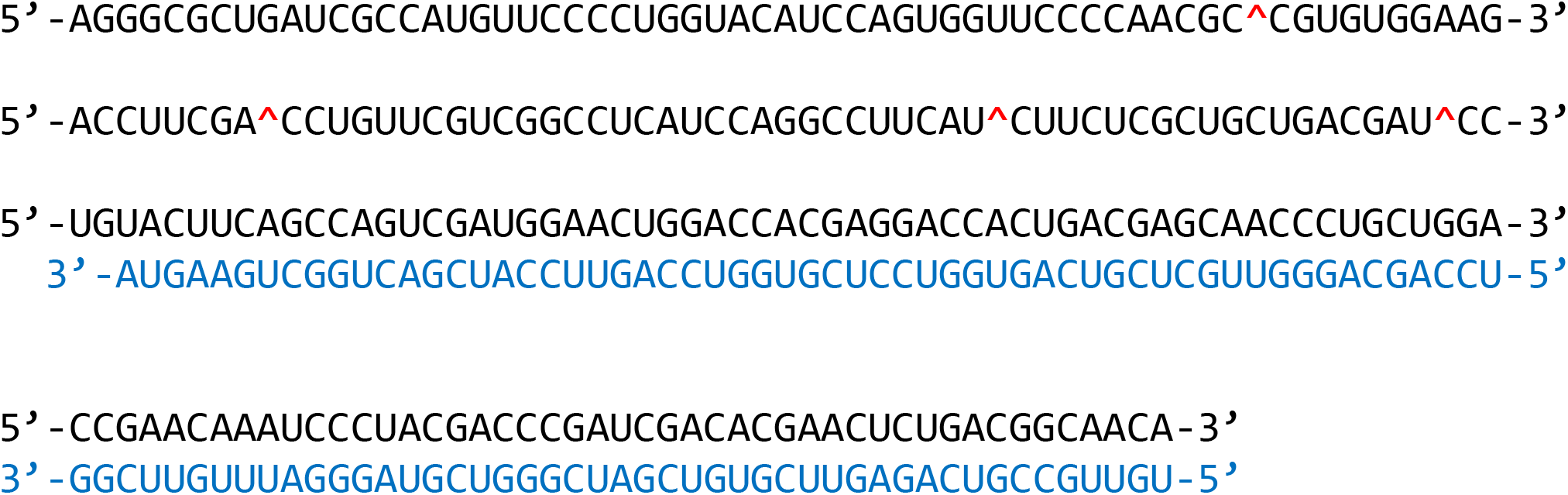
In vitro-transcribed partial duplex RNA substrate used for RNase E cleavage assays. Black font indicates the sense strand corresponding to the 3’ 159 nt of the *M. smegmatis atpB* coding sequence and 64 nt of the intergenic region between *atpB* and *atpE*. Blue font indicates an antisense strand used to block RNase E cleavage. Cleavage sites mapped in Fig. 5 are shown by red carets.

**Figure S13.**
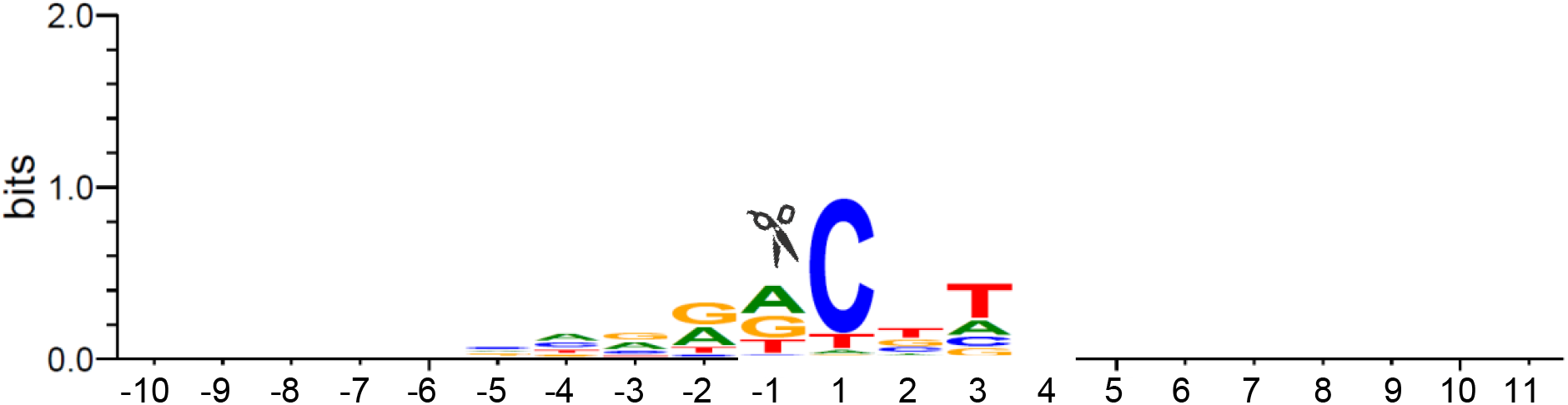
Sequence context of an expanded set of *M. tuberculosis* RNA cleavage sites. 10,795 putative cleavage sites were identified using relaxed filters as described in the methods section. A weblogo (Weblogo 3.7.12) was constructed with a background frequency of 65% G+C.

**Figure S14.**
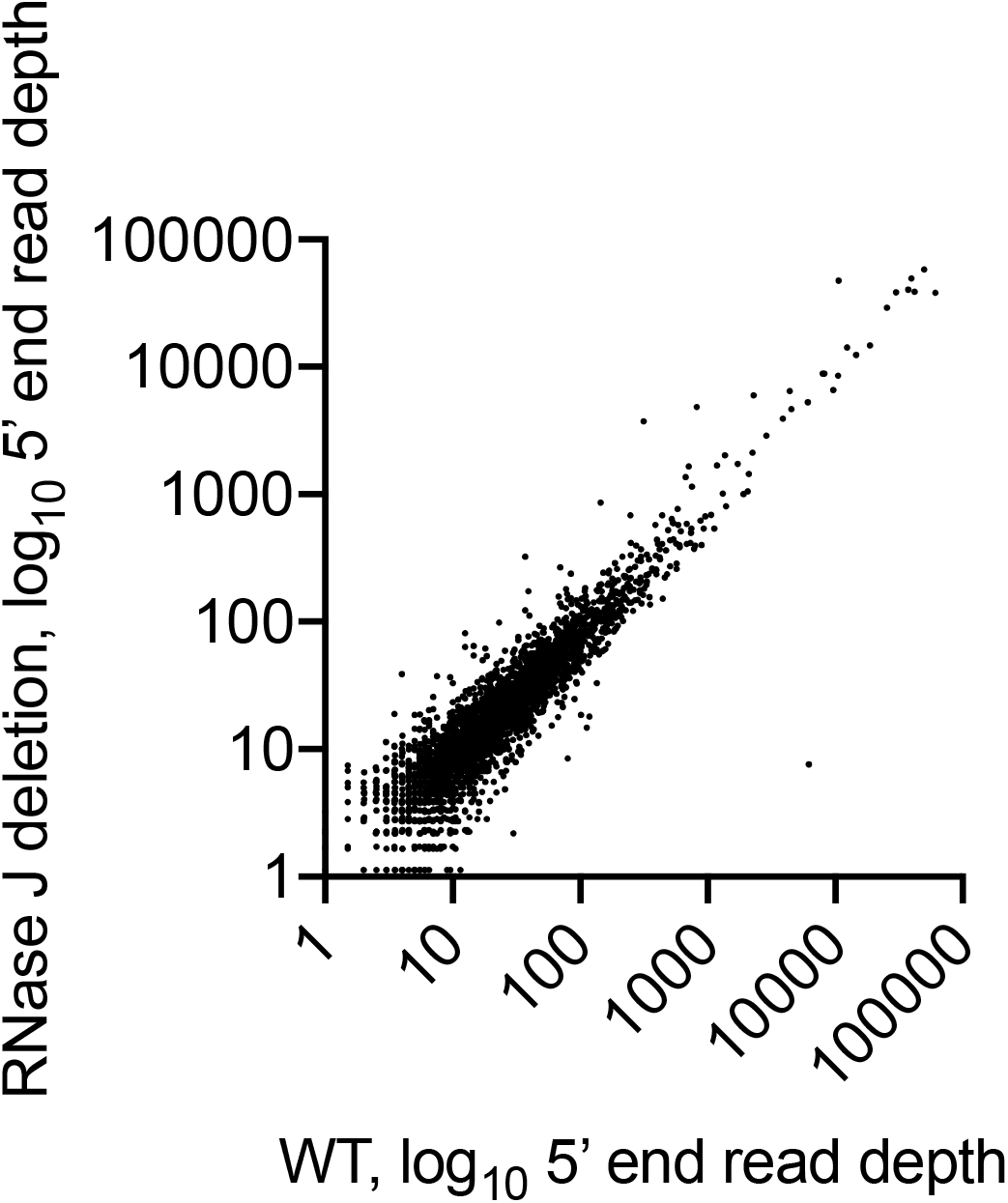
Most *M. tuberculosis* cleavage sites have similar abundance in WT H37Rv and an isogenic strain in which the gene encoding RNase J was deleted. Monophosphorylated RNA 5’ ends were mapped and quantified by adapter ligation and Illumina sequencing. Read depth for each 5’ end produced by the cleavage sites listed in Supplementary Table 5 is shown.

**Figure S15.**
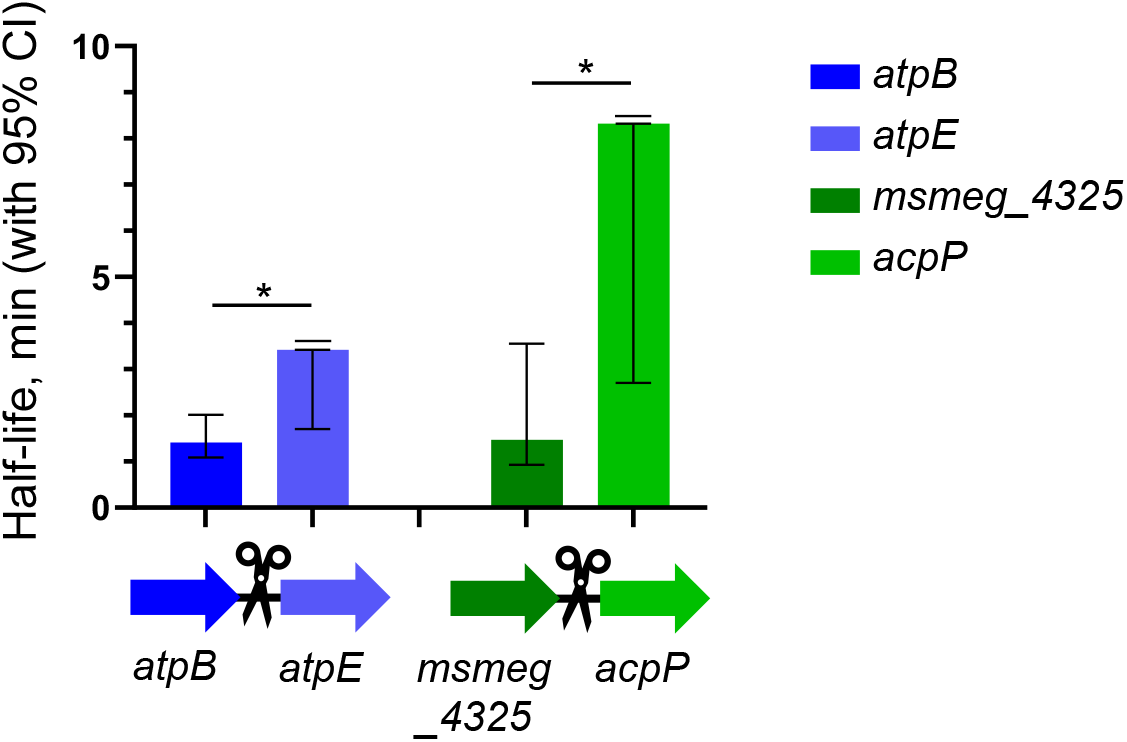
*M. smegmatis* gene pairs that appear to be co-transcribed and are bisected by cleavage sites display differential stabilities. Abundance of the four indicated transcripts was measured by qPCR 0, 4, and 8 minutes after addition of rifampicin to block transcription, and half-lives were calculated by linear regression of log_2_-transformed abundance. Error bars show the 95% confidence intervals of each half-life. The top error bar was truncated in two cases (*atpE* and *acpP*) where the upper 95% CI was infinity. *, *p* < 0.05 for comparison of the half-lives of the indicated genes by linear regression.

